# Closed-loop neurofeedback reshapes preparatory brain states to bias subsequent pain processing

**DOI:** 10.64898/2026.06.02.729491

**Authors:** Yinhua Zhang, Shuang Qiu, Qianqian Zheng, Shihao Chen, Jia Li, Xiaoyun Li, Richu Jin, Weiwei Peng

## Abstract

Pain perception is shaped not only by nociceptive input but also by neural activity preceding stimulation. Whether these preparatory states can be modified through neurofeedback, and whether such effects depend on feedback authenticity, remains unclear. We used a double-blind, sham-controlled 2 × 2 mixed factorial design, crossing active regulation versus passive observation with real versus sham EEG feedback. Across feedback groups, active training was associated with changes in preparatory EEG activity and reductions in pain intensity and unpleasantness. Compared with sham feedback, authentic EEG-contingent feedback produced greater enhancement of α power in the targeted somatosensory cortex, also evident in source estimates of the right postcentral gyrus, and selectively reduced transitions from microstate E to D. Authentic feedback was also associated with greater attenuation of laser-evoked N2 and P2 responses and a larger reduction in pain intensity, but not unpleasantness. Exploratory structural equation modeling identified indirect associations linking feedback authenticity, preparatory α and microstate changes, P2 attenuation, and pain-rating changes. Together, these findings provide experimental evidence that preparatory neural activity can be modulated through authentic neurofeedback and that this modulation is accompanied by changes in subsequent nociceptive processing, supporting a state-based account of pain.

## Introduction

Pain perception is not determined solely by nociceptive input but is strongly shaped by the brain state prior to stimulus onset ^1,2^. Converging evidence shows that pre-stimulus brain activity biases subsequent sensory processing, influencing both cortical responses and subjective pain experience ^3–5^. These preparatory states may influence the responsiveness of cortical systems to incoming nociceptive signals by modulating neural excitability and attentional engagement before stimulus onset. Among the neural mechanisms shaping these preparatory states, oscillatory activity in the α-frequency band has been consistently implicated in sensory gating and pain modulation ^6–9^. In particular, α oscillations in the primary somatosensory cortex (S1) are thought to regulate cortical excitability and filter incoming sensory input ^10,11^, with higher pre-stimulus α power often associated with attenuated pain-evoked responses and reduced pain perception ^6,12–14^. However, these correlational findings leave open whether experimentally enhancing preparatory α activity can influence subsequent pain processing and how such modulation relates to broader neural dynamics.

Closed-loop neurofeedback (NFB) provides an approach for actively modulating preparatory brain activity by translating ongoing neural signals into real-time feedback, enabling individuals to learn to regulate targeted neural activity ^15–17^. Its potential for pain modulation was first demonstrated in pioneering neurofeedback studies ^18^, and subsequent EEG-based investigations have explored the regulation of somatosensory α oscillations, although behavioral outcomes have been variable ^19–21^. An important challenge in interpreting these findings is determining whether the observed effects reflect genuine neural self-regulation or nonspecific influences associated with task engagement, attention, cognitive strategies, and expectations ^22–24^. In particular, without appropriate control conditions, improvements observed during neurofeedback cannot necessarily be attributed to the contingency between ongoing neural activity and the feedback provided. Sham-controlled designs have therefore been emphasized as essential for evaluating the specificity of neurofeedback effects ^22,25,26^. Conceptually, distinguishing the contribution of active self-regulation from that of authentic EEG-contingent feedback requires consideration of both training engagement and feedback authenticity. A factorial design combining active regulation and passive observation under real and sham feedback provides a means of separating general training-related changes from additional effects associated with authentic neural feedback. Whether these distinct contributions differentially influence preparatory neural activity and extend to subsequent nociceptive processing, however, remains unclear.

Beyond local oscillatory modulation, an important question is whether neurofeedback-related changes extend to the broader temporal organization of preparatory brain states. Although somatosensory α oscillations provide insights into local rhythmic activity and cortical excitability, they offer a limited account of how distributed neural activity evolves over time. Pain processing involves coordinated activity across multiple neural systems supporting sensory, attentional, and affective functions ^27–29^, suggesting that modulation of preparatory neural activity may involve multiple levels of brain organization. EEG microstates, characterized by brief, quasi-stable scalp electrical topographies, provide a complementary framework for examining the temporal organization and transitions of large-scale neural activity ^30,31^. Evidence from simultaneous EEG–fMRI and source-imaging studies suggests that microstate dynamics are related to distributed neural systems ^32–34^, although individual microstate classes cannot be uniquely mapped onto specific functional networks. Previous neuromodulation studies have reported changes in both oscillatory activity and microstate dynamics in association with pain modulation ^35,36^. These findings raise the possibility that targeted modulation of somatosensory α activity may be accompanied by changes in the temporal organization of preparatory brain states. Whether these neural changes are related to subsequent pain-evoked cortical responses and subjective pain perception remains unclear.

Here, we investigated whether preparatory brain states can be actively modulated through closed-loop neurofeedback and how such modulation relates to subsequent nociceptive processing and pain perception. To distinguish general training-related changes from additional effects associated with authentic feedback, we employed a double-blind, sham-controlled 2 × 2 mixed factorial design. As illustrated in Fig. 1A, participants were assigned to real- or sham-feedback groups and completed both training (active regulation) and nontraining (passive observation) conditions. At the trial level (Fig. 1B), a feedback phase preceded each nociceptive stimulus, followed by separate ratings of pain intensity and unpleasantness. During neurofeedback phase (Fig. 1C), ongoing somatosensory α power was translated into visual feedback in the real-feedback group, whereas sham feedback was generated independently of participants’ ongoing EEG activity. We hypothesized that authentic feedback would enhance preparatory somatosensory α regulation and produce greater training-related modulation of subsequent pain responses. By integrating EEG oscillations, microstate dynamics, pain-evoked cortical responses, and subjective perception, this study aimed to extend the investigation of preparatory brain states from their predictive associations with pain toward their experimental modulation and relationship with subsequent pain processing.

**Figure 1.**
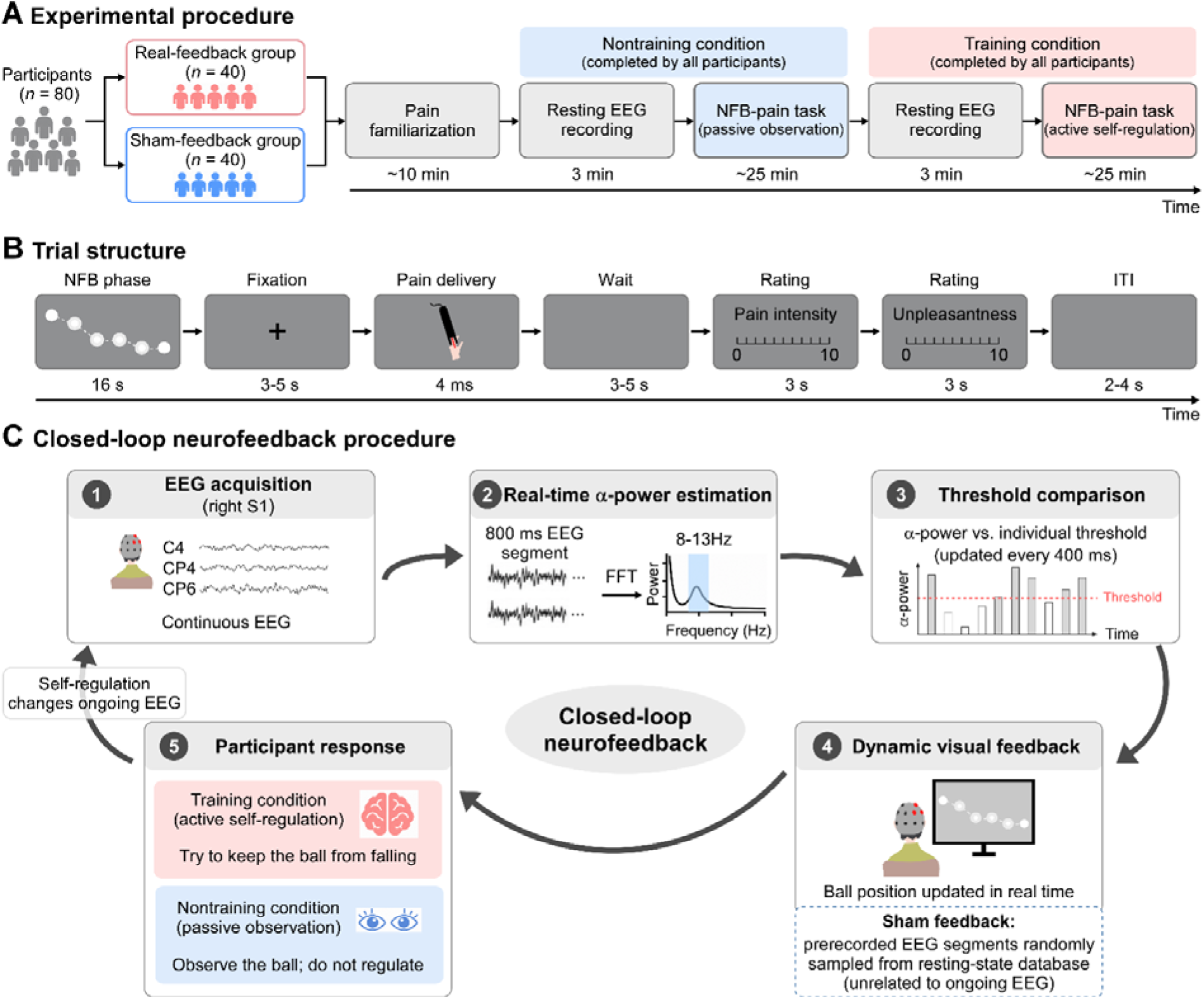
Experimental design and closed-loop neurofeedback procedure. ***(A) Experimental procedure.*** Participants were assigned to either the real-feedback or sham-feedback group (initial *n* = 40 per group). Following pain familiarization (∼10 min), all participants completed both nontraining (passive observation) and training (active self-regulation) conditions. Each condition began with a 3-min resting-state EEG recording, followed by a neurofeedback–pain task (∼25 min). Thus, Training was manipulated within participants, whereas Feedback authenticity was manipulated between participants. ***(B) Trial structure.*** Each trial consisted of a 16-s neurofeedback phase, a fixation interval (3–5 s), a 4-ms noxious laser stimulus, a post-stimulus interval (3–5 s), separate ratings of pain intensity and unpleasantness (0–10; 3 s each), and an intertrial interval (2–4 s). ***(C) Closed-loop neurofeedback procedure.*** In the real-feedback group, ongoing EEG activity was recorded from electrodes C4, CP4, and CP6 over the right somatosensory cortex (target S1). Absolute α power (8–13 Hz) was estimated from 800-ms EEG windows, updated every 400 ms, and averaged across the three electrodes. The estimated α power was compared with an individualized reference threshold derived from the 3-min resting-state EEG recording immediately preceding the corresponding condition, and the comparison determined the position of the visual feedback ball. In the sham-feedback group, both the feedback signal and reference threshold were derived from prerecorded EEG data obtained from independent donors and processed using the same procedure; thus, the feedback was unrelated to participants’ ongoing EEG activity. During training, participants actively attempted to keep the ball from falling, whereas during nontraining, they passively observed its movement without attempting self-regulation.

## Results

As shown in Table 1, the real- and sham-feedback groups did not differ in demographic variables (age, sex) or baseline psychological measures (all *p* > 0.38), indicating good comparability prior to the intervention. Blinding was effective, as participants’ beliefs about their assigned condition did not differ between groups (χ^2^*(1)* = 0.83, *p* = 0.36).

**Table 1.** Demographic and psychological characteristics of participants.

|  | **Real (*n* = 38)** | **Sham (*n* = 36)** | **Statistics** |
|---|---|---|---|
| **Sex (Male/Female)** | 20/18 | 21/15 | *χ<sup>2</sup><sub>(1)</sub>* = 0.24 |
| **Age (years)** | 20.16 ± 0.39 | 20.25 ± 0.35 | *t*<sub>72</sub> = −0.18 |
| **Treatment blindness (Yes/No)** | 28/10 | 23/13 | *χ<sup>2</sup><sub>(1)</sub>* = 0.83 |
| **Pain catastrophizing** | 22.42 ± 1.85 | 23.28 ± 1.65 | *t*<sub>72</sub> = −0.34 |
| **Pain sensitivity** | 66.89 ± 2.45 | 69.53 ± 3.34 | *t*<sub>72</sub> = −0.64 |
| **Fear of pain** | 99.97 ± 2.31 | 102.86 ± 2.35 | *t*<sub>72</sub> = −0.88 |
| **Pain anxiety symptoms** | 46.68 ± 2.96 | 46.86 ± 2.42 | *t*<sub>72</sub> = −0.05 |
| **Pain resistance** | 50.47 ± 1.97 | 48.92 ± 1.84 | *t*<sub>72</sub> = 0.58 |
| **State anxiety** | 40.13 ± 1.74 | 41.28 ± 1.58 | *t*<sub>72</sub> = −0.49 |
| **Trait anxiety** | 43.34 ± 1.50 | 43.78 ± 1.36 | *t*<sub>72</sub> = −0.21 |
| **Emotion regulation** |  |  |  |
| Frequency of reappraisal | 31.55 ± 0.80 | 30.89 ± 0.89 | *t*<sub>72</sub> = 0.56 |
| Frequency of suppression | 14.97 ± 0.87 | 15.14 ± 0.82 | *t*<sub>72</sub> = −0.14 |

**Note.** Data are presented as mean ± SEM. Between-group comparisons (real vs. sham) were conducted using independent-sample *t*-tests for continuous variables and χ² tests for categorical variables.

During the experiment, participants were not instructed to use specific regulation strategies but were asked to report the strategies they had adopted during training. Memory and imagery were most frequently reported (86.49%), followed by attentional focus (52.70%), somatic or physiological regulation (47.30%), and verbal or numerical cognitive strategies (45.95%). The strategies were not mutually exclusive, as participants could report multiple approaches (Supplementary Table S1).

### Neurofeedback effects on EEG oscillations

Grand-average power spectra and training-related changes are shown in Fig. 2A. To account for multiple comparisons, false discovery rate (FDR) correction was performed separately across the five frequency bands within each region of interest (ROI) ^37^. Corrected *p* values are reported as *p_FDR_*. Two-way mixed-design ANOVAs revealed significant main effects of Training across multiple frequency bands (Supplementary Table S2). Relative to nontraining, active training increased θ- and α-power in the target and nontarget somatosensory and frontal regions, while decreasing β- and γ-power across regions and δ-power in the nontarget somatosensory and parietal regions (all *F*_1,72_ > 4.52, *p_FDR_* < 0.05, η_p_^2^> 0.05). These findings indicate widespread oscillatory changes associated with active training, irrespective of feedback authenticity.

**Figure 2.**
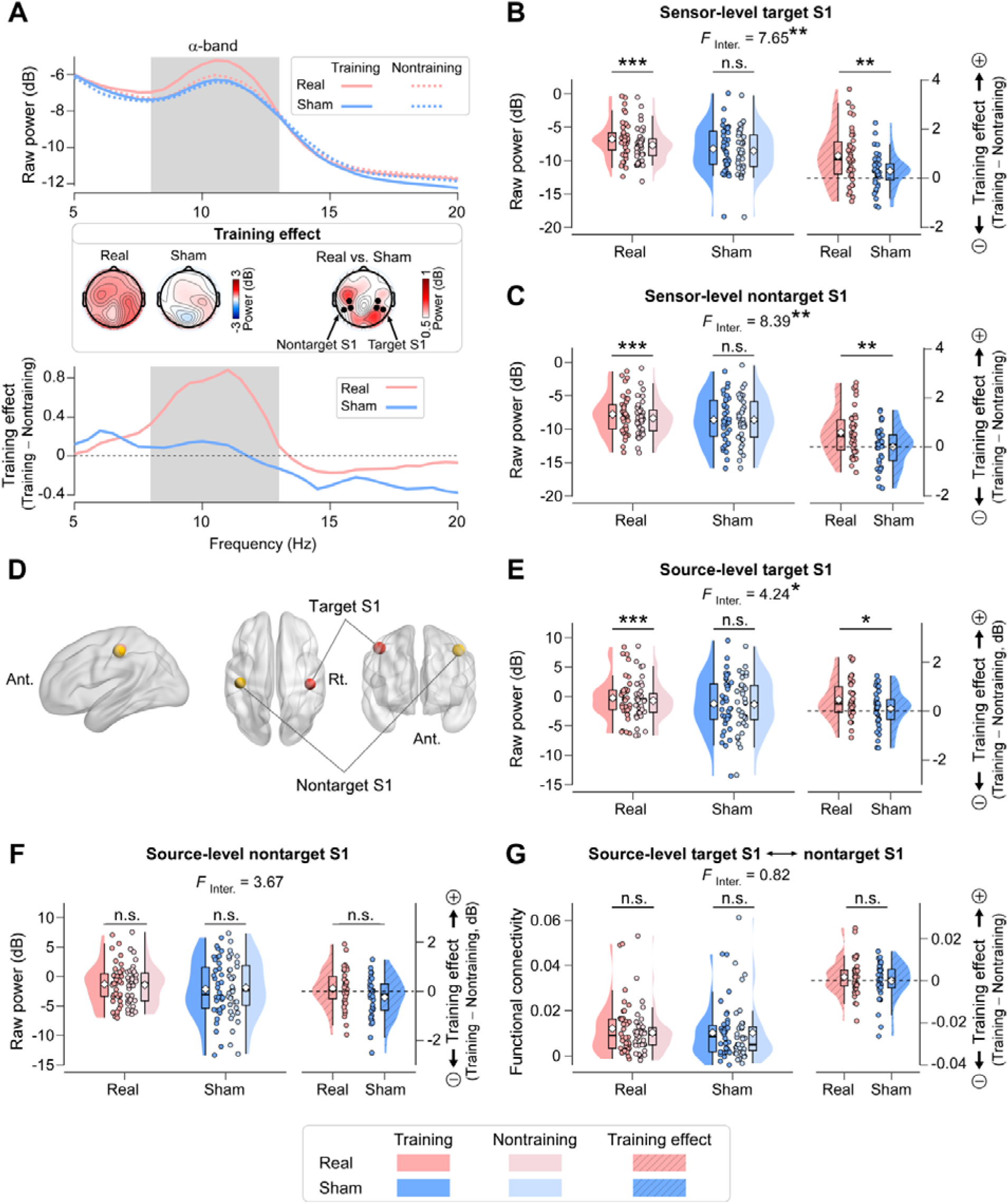
Neurofeedback-related modulation of somatosensory α oscillations and interhemispheric functional connectivity at sensor and source levels. ***(A)*** *EEG power spectra.* Grand-average power spectral density (5–20 Hz) from the target somatosensory electrodes (C4, CP4, and CP6) is shown for training and nontraining conditions in the real- and sham-feedback groups. The lower panel shows training-related changes (Training − Nontraining). Scalp maps illustrate the corresponding changes in α power (8–13 Hz) and their between-group differences. *(B–C) Sensor-level* α *power.* α power in the target (B) and nontarget (C) somatosensory regions is shown for both feedback groups. Significant Training × Feedback authenticity interactions were observed in both regions, with greater training-related α enhancement under real than sham feedback. *(D) Source-space regions of interest.* The bilateral postcentral gyri were anatomically defined as the source-space counterparts of the target (right S1) and nontarget (left S1) somatosensory regions. *(E–F) Source-level* α *power.* Source-reconstructed α power in the right (E) and left (F) postcentral gyri is shown for training and nontraining conditions in both feedback groups. A significant Training × Feedback authenticity interaction was observed in the right postcentral gyrus, with a training-related increase under real feedback but not sham feedback. The corresponding interaction in the left postcentral gyrus did not reach significance. *(G) Interhemispheric functional connectivity.* α-band functional connectivity between the bilateral somatosensory regions is shown for training and nontraining conditions in both feedback groups. No significant Training × Feedback authenticity interaction was observed. For panels B, C, E, F, and G, the right-hand plots show within-participant training-related changes (Training − Nontraining). Violin plots show data distributions; box plots indicate medians and interquartile ranges; diamonds indicate means; dots represent individual participants. Significance levels: **\***, *p* < 0.05; **\*\***, *p* < 0.01; **\*\*\***, *p* < 0.001; n.s., not significant. Abbreviations: Inter., Training × Feedback authenticity interactions; “+” indicates a greater training effect, whereas “−” indicates a smaller training effect.

Critically, significant Training × Feedback authenticity interactions were observed for α-power in both the target (*F*_1, 72_ = 7.65, *p_FDR_* = 0.04, η_p_^2^= 0.10; Fig. 2B) and nontarget somatosensory regions (*F*_1, 72_ = 8.39, *p_FDR_* = 0.02, η_p_^2^ = 0.10; Fig. 2C). Post hoc comparisons revealed significant training-related α-power increases in the real-feedback group (both *p* < 0.001), whereas the corresponding changes in the sham-feedback group were not significant (*p* = 0.06 and 0.99, respectively). Accordingly, training-related α enhancement was significantly greater under real than sham feedback in both regions (*p* = 0.007 and 0.005, respectively). These findings indicate that authentic EEG-contingent feedback enhanced somatosensory α regulation beyond the changes associated with active training under sham feedback.

To further characterize the temporal dynamics of α regulation, we examined target-S1 α power across different windows of the 16-s feedback period. The Training × Feedback authenticity interaction remained significant during the first 8 s (*F*_1, 72_ = 5.73, *p* = 0.02, η_p_^2^ = 0.07), last 8 s (*F*_1, 72_ = 5.99, *p* = 0.02, η_p_^2^ = 0.08), and final 2 s (*F*_1, 72_ = 6.71, *p* = 0.01, η_p_^2^ = 0.09), indicating that feedback-dependent α modulation was detectable throughout the feedback period (Supplementary Fig. S1A). However, during the final 2 s of the subsequent fixation interval immediately preceding laser stimulation, the Training × Feedback authenticity interaction was not significant (*F*_1, 72_ = 0.10, *p* = 0.75, η_p_^2^= 0.001; Supplementary Fig. S1B), despite a significant main effect of Training (*F*_1, 72_ = 75.45, *p* < 0.001, η_p_^2^= 0.51). Thus, the additional α-regulation effect associated with authentic feedback was consistently observed during the feedback period but was not detectable immediately before nociceptive stimulation.

To further validate these findings at the cortical source level, we reconstructed EEG activity using a linearly constrained minimum variance (LCMV) beamformer and extracted α power from the bilateral postcentral gyri (Fig. 2D). Consistent with the sensor-level results, source α power in the right postcentral gyrus (target S1) showed a significant Training × Feedback authenticity interaction (*F*_1, 72_ = 4.24, *p* = 0.04, η_p_^2^ = 0.06; Fig. 2E). Follow-up comparisons revealed a significant training-related increase in the real-feedback group (*p* < 0.001), but not in the sham-feedback group (*p* = 0.44). In contrast, the corresponding interaction in the left postcentral gyrus (nontarget S1) did not reach statistical significance (*F*_1, 72_ = 3.67, *p* = 0.06, η_p_^2^ = 0.05; Fig. 2F). Thus, the source-space analysis provided additional support for feedback-dependent α modulation in the targeted right somatosensory cortex, whereas the evidence for a corresponding effect in the left hemisphere was less conclusive.

We additionally examined interhemispheric α-band connectivity between the bilateral postcentral gyri using the debiased weighted phase-lag index (dwPLI). This source-space analysis did not reveal a significant Training × Feedback authenticity interaction (*F*_1, 72_ = 0.82, *p* = 0.37, η_p_^2^ = 0.01; Fig. 2G). Therefore, the original sensor-level connectivity finding was not supported by the more conservative source-space analysis, and no connectivity-based mechanism was inferred.

### Neurofeedback effects on EEG microstates

Topographic maps of the five canonical microstate classes exhibited high spatial consistency across groups (all *r*_60_ > 0.95, *p* < 0.001; Fig. 3A; Supplementary Fig. S2), with a mean global explained variance of 78.76%, indicating comparable microstate structure across conditions.

**Figure 3.**
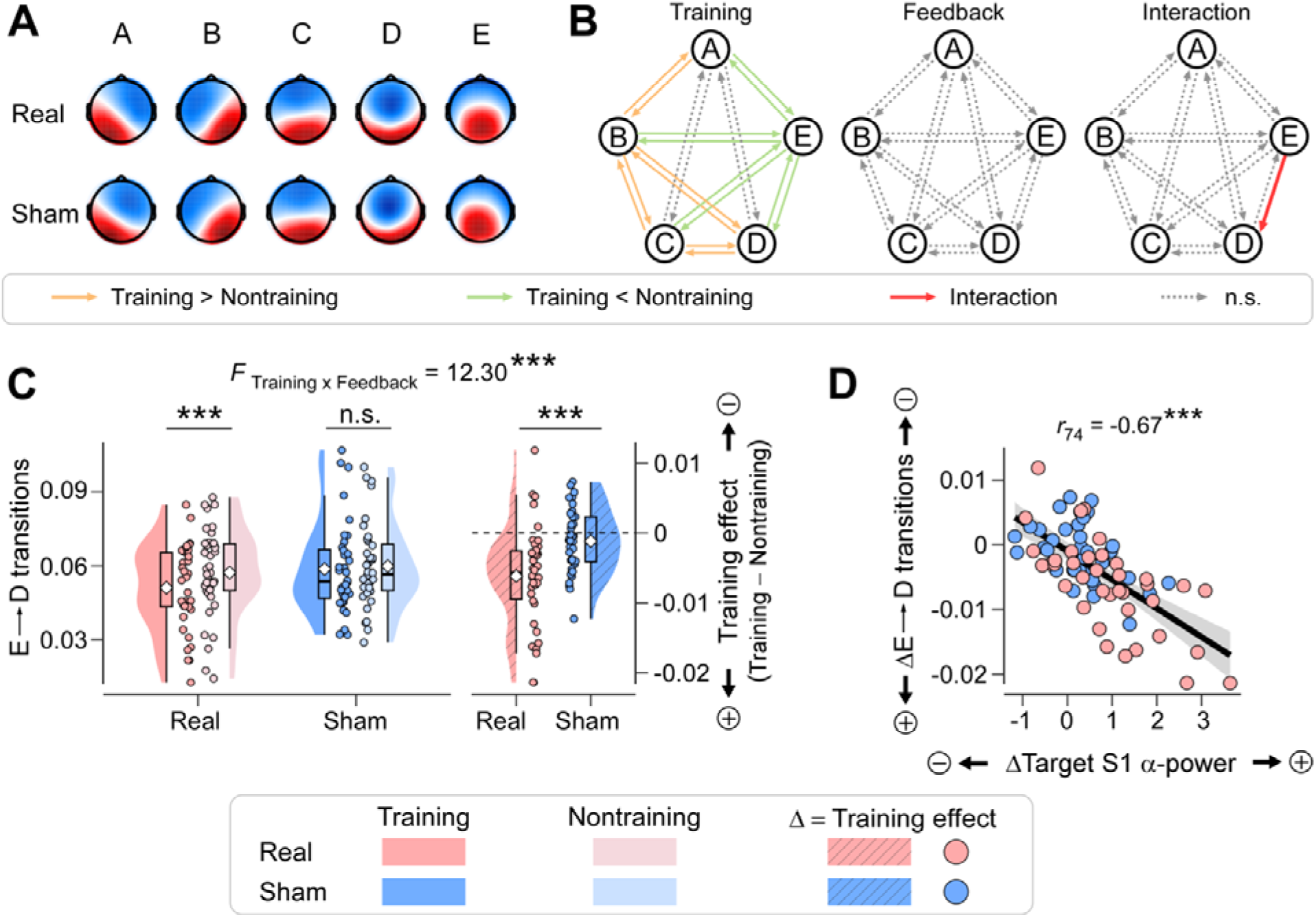
Neurofeedback-related changes in preparatory EEG microstate dynamics. ***(A) Spatial configurations of EEG microstates.*** Five canonical microstate classes (A–E) were identified across the real- and sham-feedback groups, showing highly consistent spatial configurations. ***(B) Training- and feedback-related modulation of microstate transitions.*** The left, middle, and right panels show the statistical effects of Training, Feedback authenticity, and their interaction, respectively, across all 20 directed transitions between the five microstate classes. Training produced widespread changes in transition probabilities, whereas only the E→D transition showed a significant Training × Feedback authenticity interaction after FDR correction. ***(C) E*** → ***D transition probability.*** Training significantly reduced E→D transitions in the real-feedback group, but not in the sham-feedback group. The magnitude of this reduction was significantly greater under real than sham feedback, consistent with the significant Training × Feedback authenticity interaction. Violin plots show data distributions; box plots indicate medians and interquartile ranges; diamonds indicate means; dots represent individual participants. ***(D) Association between E***→***D transition probability and somatosensory*** α ***modulation.*** Across participants from both feedback groups, greater training-related reductions in E→D transition probability were associated with larger increases in target-S1 α power (Δ = Training − Nontraining). Dots represent individual participants (red = real feedback; blue = sham feedback). The solid line indicates the linear fit, with shaded areas representing the 95% confidence interval. Significance levels: **\*\*\***, *p* < 0.001; ns., not significant. Abbreviations: “+” indicates a greater training effect, whereas “−” indicates a smaller training effect.

Two-way mixed-design ANOVAs revealed significant main effects of Training on several microstate temporal characteristics (Supplementary Table S3). To account for multiple comparisons, FDR correction was applied across the 15 temporal characteristics (mean duration, occurrence, and time coverage across five microstates). Compared with nontraining, training increased the mean duration of microstates A, B, and D and the time coverage of microstate B (all *F*_1,72_ > 6.70, *p_FDR_* < 0.03, η_p_^2^ > 0.08), while reducing the duration, occurrence, and time coverage of microstate E (all *F*_1,72_ > 8.53, *p_FDR_* < 0.02, η_p_^2^ > 0.10). No significant Training × Feedback authenticity interactions were observed for any temporal characteristic after FDR correction (all *p_FDR_* > 0.26; Supplementary Fig. S3A−C), indicating comparable training-related changes across feedback groups.

Analyses of microstate transition probabilities are summarized in Supplementary Table S4. FDR correction was applied across the 20 directed microstate transition probabilities to account for multiple comparisons. Compared with the nontraining condition, training facilitated transitions among microstates B↔A, B↔C, B↔D, and C↔D (all *F*_1,72_ > 6.74, *p_FDR_*< 0.02, η_p_^2^> 0.08), while suppressing transitions involving microstate E (E↔A, E↔B, E↔C, E↔D; all *F*_1,72_ > 4.49, *p_FDR_* < 0.05, η_p_^2^ > 0.05; Fig. 3B). Among the 20 directed transitions, only E→D showed a significant Training × Feedback authenticity interaction after FDR correction (*F*_1,72_ = 12.30, *p_FDR_* = 0.02, η_p_^2^ = 0.15; Supplementary Fig. S3D). Post hoc analyses showed that training robustly reduced E→D transitions in the real-feedback group (*p* < 0.001), but not in the sham-feedback group (*p* = 0.26), with a significantly stronger reduction under real feedback (*p* < 0.001; Fig. 3C). Moreover, across participants from both feedback groups, training-related reductions in E→D transitions were negatively correlated with increases in target-region α power (*r*_74_ = −0.67, *p* < 0.001; Fig. 3D), directly linking microstate dynamics to oscillatory modulation.

These findings indicate that active training was accompanied by broad changes in microstate dynamics, whereas authentic feedback was associated with a selective additional reduction in E→D transitions, which was correlated with somatosensory α modulation.

### Neurofeedback effects on pain ratings and evoked potentials

Pain ratings and training-induced changes are illustrated in Fig. 4A and Fig. 4B. Two-way mixed-design ANOVAs revealed robust main effects of training on both pain intensity (*F*_1,72_ = 68.32, *p* < 0.001, η_p_^2^ = 0.49) and pain unpleasantness (*F*_1,72_ = 53.66, *p* < 0.001, η_p_^2^ = 0.43), indicating lower pain ratings during training relative to nontraining across groups. Main effects of feedback authenticity further showed generally lower ratings in the real-feedback group for both intensity (*F*_1,72_ = 5.66, *p* = 0.02, η_p_^2^ = 0.07) and unpleasantness (*F*_1,72_ = 4.41, *p* = 0.04, η_p_^2^ = 0.06).

**Figure 4.**
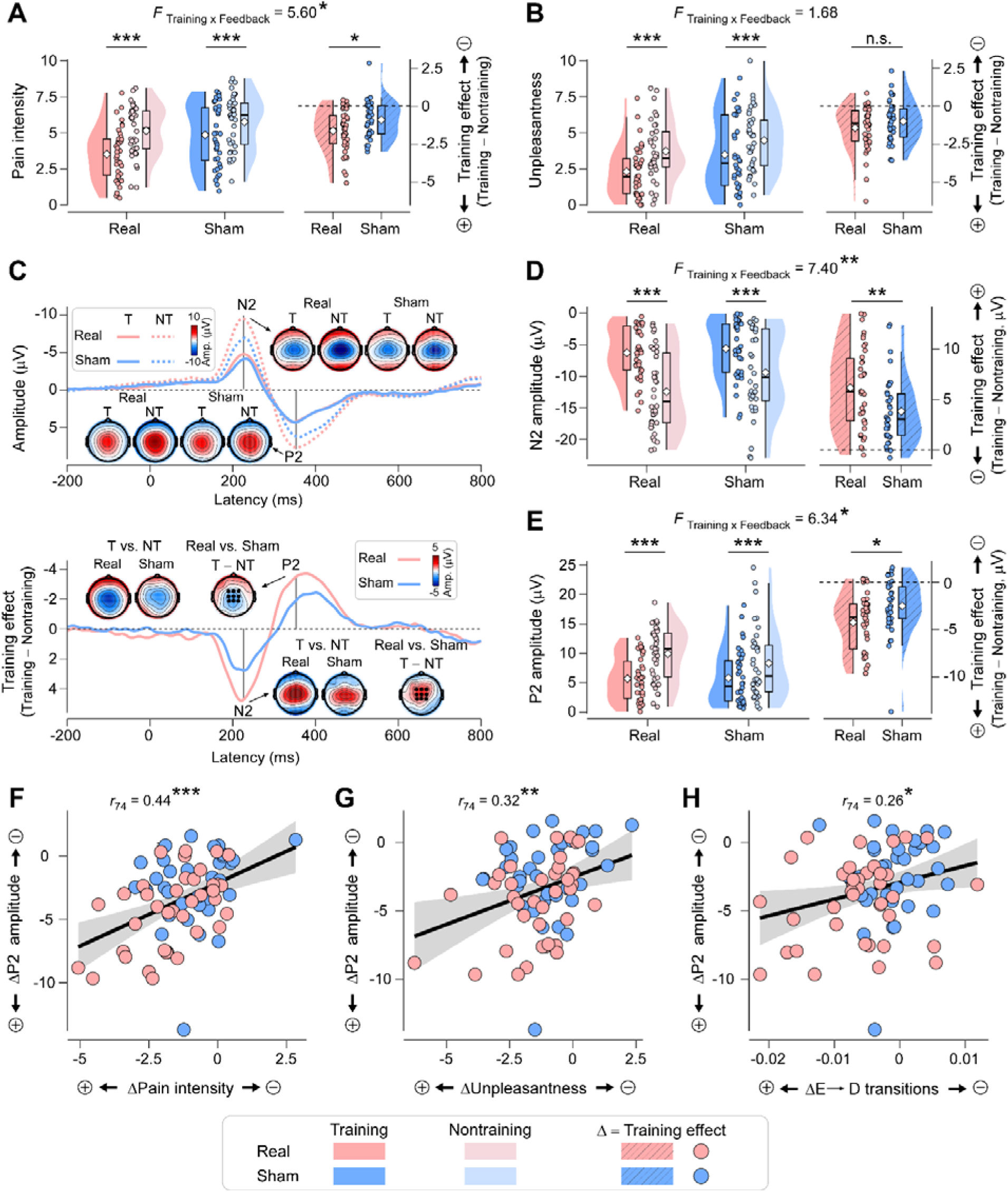
Effects of neurofeedback on subjective pain perception and pain-evoked cortical responses. ***(A–B) Pain ratings.*** Subjective ratings of pain intensity (A) and unpleasantness (B) elicited by noxious laser stimulation are shown for training and nontraining conditions under real (red) and sham (blue) feedback. Training reduced both pain intensity and unpleasantness across groups. A significant Training × Feedback authenticity interaction was observed for pain intensity, with a greater reduction under real than sham feedback, whereas no significant interaction was observed for unpleasantness. ***(C) Laser-evoked potentials (LEPs).*** Upper panel: Grand-average LEP waveforms for training and nontraining conditions in the real- and sham-feedback groups, together with scalp topographies of the N2 and P2 components. Lower panel: Difference waveforms (Training − Nontraining) and corresponding scalp topographies illustrating training-related changes in N2 and P2 responses. ***(D–E) N2 and P2 amplitudes.*** Training significantly reduced N2 and P2 amplitudes for both real and sham neurofeedback groups. The training effects on reducing these two components (training − nontraining) were significantly greater for real than sham feedback. Violin plots show data distributions; box plots indicate medians and interquartile ranges; diamonds indicate means; dots represent individual participants. ***(F–H) Associations between pain-evoked responses, pain ratings, and preparatory brain dynamics.*** Across participants, greater training-related reductions in P2 amplitude were positively associated with decreases in pain intensity and unpleasantness, as well as with reduced E → D microstate transition probability. Each dot represents an individual participant (red = real feedback; blue = sham feedback); black lines indicate linear fits, with gray shading denoting 95% confidence intervals. Significance levels: **\***, *p* < 0.05; **\*\***, *p* < 0.01; **\*\*\***, *p* < 0.001; n.s., not significant. Abbreviations: T, training; NT, nontraining; “+” indicates a greater training effect, whereas “−” indicates a smaller training effect.

Importantly, a significant training × feedback interaction was observed for pain intensity (*F*_1,72_ = 5.60, *p* = 0.02, η_p_^2^ = 0.07), but not for unpleasantness (*F*_1,72_ = 1.68, *p* = 0.20, η_p_^2^= 0.02). Post hoc analyses confirmed that pain intensity decreased significantly in both groups (all *p* < 0.001), with a larger reduction under real feedback (*p* = 0.02; Fig. 4A). Pain unpleasantness decreased comparably in both groups (all *p* < 0.001), with no group difference (*p* = 0.20; Fig. 4B). After adjustment for positive-feedback proportion in linear mixed-effects models, the Training × Feedback authenticity interaction remained significant for pain intensity *(F* = 5.26, *p* = 0.02), but not for unpleasantness (*F* = 1.13, *p* = 0.29). These findings indicate that active training was associated with reductions in both dimensions of pain, while authentic feedback conferred an additional benefit for pain intensity that was not explained by differences in positive-feedback proportion.

Grand-average LEP waveforms and training-induced changes are shown in Fig. 4C. Consistent with behavioral effects, neurofeedback significantly modulated cortical pain responses. Robust main effects of training were observed for both the N2 (*F*_1,72_ = 135.05, *p* < 0.001, η_p_^2^ = 0.65) and P2 components (*F*_1,72_ = 96.70, *p* < 0.001, η_p_^2^ = 0.57), reflecting reduced amplitudes during training relative to nontraining. As shown in Fig. 4D and Fig. 4E, significant training × feedback interactions were also detected for both N2 (*F*_1,72_ = 7.40, *p* = 0.008, η_p_^2^ = 0.09) and P2 (*F*_1,72_ = 6.34, *p* = 0.01, η_p_^2^ = 0.08). Post hoc tests showed amplitude reductions in both groups (all *p* < 0.001), with significantly larger decreases under real feedback (all *p* < 0.02).

Finally, reductions in LEP amplitudes were associated with behavioral pain relief. Training-related decreases in N2 and P2 amplitudes correlated with reductions in pain intensity (N2: *r*_74_ = −0.37, *p* = 0.001; P2: *r*_74_ = 0.44, *p* < 0.001; Fig. 4F) and unpleasantness (N2: *r*_74_ = −0.25, *p* = 0.03; P2: *r*_74_ = 0.32, *p* = 0.005; Fig. 4G). Moreover, greater P2 attenuation was associated with stronger suppression of E→D microstate transitions (*r*_74_ = 0.26, *p* = 0.02; Fig. 4H). Together, these findings indicate that authentic feedback enhanced the analgesic efficacy of neurofeedback by jointly attenuating pain-evoked cortical responses and reshaping large-scale brain state dynamics.

To assess potential within-session temporal effects, we divided the 30 trials into early and late phases and incorporated Phase into the mixed-design ANOVAs. Significant Phase effects were observed for pain ratings, N2/P2 amplitudes, and somatosensory α power, indicating changes in behavioral and neural responses over time. Importantly, the Training × Feedback authenticity interactions remained significant for pain intensity, N2/P2 amplitudes, and bilateral somatosensory α power (all *p* ≤ 0.02), with no significant moderation by Phase (all three-way interaction *p* > 0.35; Supplementary Table S5).

### Associations among preparatory neural dynamics and subsequent pain processing

To examine whether somatosensory α modulation was associated with pain reduction, we first assessed the relationship between training-related changes in target-S1 α power and pain intensity, which was not significant across participants (*r*_74_ = −0.09, *p* = 0.44; Supplementary Table S6). Trial-level mixed-effects analyses revealed that higher α power was associated with lower subsequent pain intensity (β = −0.15, *p* = 0.006; Supplementary Table S7). However, this association was attenuated during training (α × Training: β = 0.15, *p* = 0.049) and was not significantly moderated by feedback authenticity (α × Feedback authenticity: *p* = 0.96; α × Training × Feedback authenticity: *p* = 0.62). Moreover, the α coefficient was no longer significant after accounting for concurrent activity in other frequency bands (β = −0.03, *p* = 0.69; Supplementary Table S8).

To further examine the relationships among feedback authenticity, preparatory neural modulation, and subsequent pain processing, we constructed a structural equation model (SEM) incorporating training-related changes (Training − Nontraining) in somatosensory α power, E→D microstate transition probability, pain-evoked P2 amplitude, and pain ratings, with Feedback authenticity (real vs. sham) specified as the exogenous variable. The model showed excellent fit (*n* = 74, χ*^2^* = 1.11, root mean square error of approximation [RMSEA] < 0.001, root mean squared residual [RMR] = 0.03, goodness-of-fit index [GFI] > 0.99), indicating that the hypothesized structure adequately captured the data.

As illustrated in Fig. 5, three significant indirect pathways were identified. First, Feedback authenticity was associated with training-related reductions in E→D transition probability, which were further associated with attenuated P2 responses and lower pain ratings (indirect effect = −0.07, 95% CI [−0.25, −0.01]). Second, Feedback authenticity was associated with greater P2 attenuation, which was related to reduced pain ratings (indirect effect = −0.26, 95% CI [−0.66, −0.02]). Finally, a sequential indirect pathway linked Feedback authenticity to increased somatosensory α power, reduced E→D transitions, attenuated P2 responses, and lower pain ratings (indirect effect = −0.07, 95% CI [−0.23, −0.01]).

**Figure 5.**
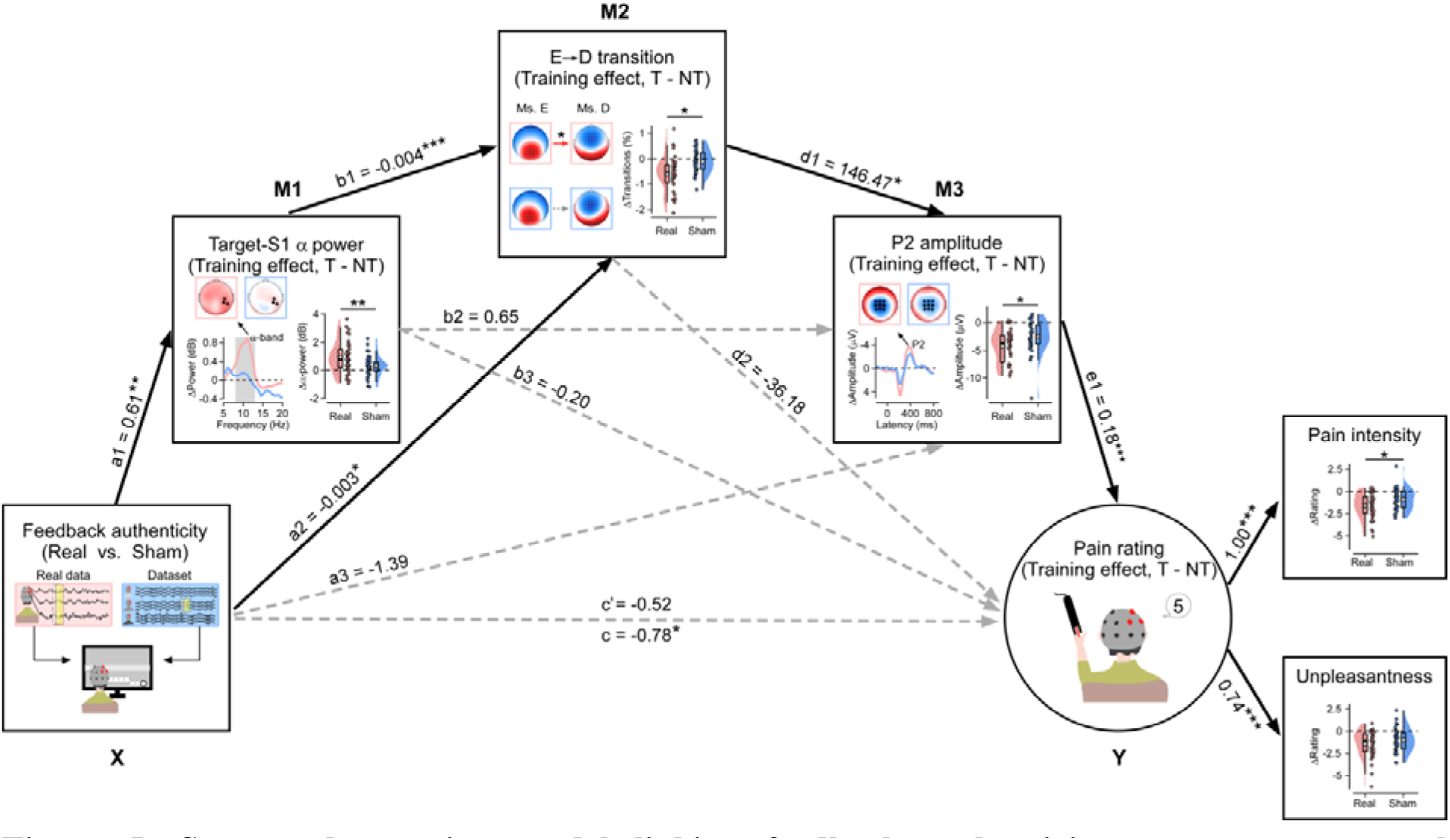
Structural equation model linking feedback authenticity, preparatory neural modulation, and pain regulation. The structural equation model examined the hypothesized direct and indirect associations linking feedback authenticity, preparatory neural changes, and subsequent pain-related outcomes. Feedback authenticity was specified as the exogenous variable (0 = sham; 1 = real). All neural and behavioral variables represent training-related changes (Δ = Training − Nontraining). M1 represents the change in target-S1 α power (Δα power); M2 represents the change in E→D microstate transition probability (ΔE→D); and M3 represents the change in pain-evoked P2 amplitude (ΔP2). The outcome variable (Y) is a latent pain-rating change factor derived from training-related changes in pain intensity and unpleasantness ratings (Δpain ratings). Solid arrows indicate statistically significant paths (*p* < 0.05), whereas dashed arrows indicate nonsignificant paths. Nonstandardized regression coefficients are shown alongside the corresponding paths. Significance levels: **\***, *p* < 0.05; **\*\***, *p* < 0.01; **\*\*\***, *p* < 0.001.

These results reveal associations among preparatory α activity, microstate dynamics, pain-evoked cortical responses, and subjective pain. Although the sequential pathway provides a potential framework for understanding how preparatory neural modulation relates to subsequent pain processing, the direct causal contribution of α enhancement to analgesia remains to be established.

## Discussion

In this study, we demonstrate that closed-loop neurofeedback can modulate preparatory brain states and influence subsequent pain processing. Across feedback groups, active training was associated with broad changes in preparatory neural activity, attenuated pain-evoked cortical responses, and reduced subjective pain. Critically, authentic EEG-contingent feedback produced greater training-related enhancement of somatosensory α oscillations, accompanied by larger reductions in pain intensity and laser-evoked N2 and P2 amplitudes compared with sham feedback. At the level of large-scale brain dynamics, authentic feedback was also associated with a selective reduction in E→D microstate transitions, suggesting changes in the temporal organization of preparatory neural states. Integrating neural and behavioral measures further revealed associations among somatosensory α modulation, microstate dynamics, pain-evoked cortical responses, and subjective pain, suggesting potential relationships across different stages of pain processing. Together, these findings extend our understanding of preparatory brain states from their predictive associations with pain toward their experimental regulation, highlighting the contribution of authentic neural feedback to preparatory neural dynamics and subsequent pain processing.

At the oscillatory level, active training was associated with widespread changes in EEG power, including an overall increase in somatosensory α activity across feedback groups. This general effect may partly reflect processes accompanying active engagement, such as attentional allocation and cognitive strategy use, consistent with evidence that somatosensory α oscillations are sensitive to top-down attentional demands ^20,38,39^. Beyond these general training effects, authentic EEG-contingent feedback produced significantly greater α enhancement in the targeted somatosensory cortex than sham feedback, highlighting the additional contribution of reliable feedback contingency to neural self-regulation. This finding is consistent with neurofeedback accounts proposing that veridical information about ongoing neural activity enables individuals to evaluate and adjust their regulatory strategies ^16,17,40^. Notably, the feedback-dependent α enhancement remained detectable throughout the 16-s neurofeedback period, including its final 2 s. However, it did not progressively increase across trials and was no longer detectable immediately before nociceptive stimulation. Thus, authentic feedback facilitated α regulation during active feedback engagement, but the present findings do not establish progressive skill acquisition or sustained modulation after feedback termination.

The spatial distribution of α modulation further informs its neural specificity. At the sensor level, authentic feedback enhanced α power in both targeted and contralateral somatosensory regions, suggesting that the modulation extended beyond the electrodes used for feedback. Source-space analyses confirmed a significant effect in the right postcentral gyrus, whereas the corresponding effect in the left postcentral gyrus showed a similar but nonsignificant trend. Importantly, no significant enhancement of interhemispheric α-band functional connectivity was observed, indicating that the bilateral sensor-level effects do not necessarily reflect increased interhemispheric synchronization. Overall, the strongest evidence for feedback-dependent α modulation was localized to the targeted S1, with broader spatial expression at the sensor level. Functionally, this modulation is relevant to established accounts of α oscillations as a mechanism for regulating cortical excitability and sensory information processing through inhibitory gating ^10,41^. Our findings extend this literature by demonstrating that authentic EEG-contingent feedback can be used to modulate preparatory somatosensory α activity, although whether this modulation directly alters sensory gating or contributes causally to subsequent analgesia remains to be established.

Beyond local oscillatory modulation, our findings suggest that neurofeedback training also influences the temporal organization of preparatory brain states, as reflected in EEG microstate dynamics. EEG microstates capture brief, quasi-stable scalp electrical configurations and their transitions, providing a complementary perspective on large-scale neural dynamics beyond regional oscillatory power ^30,31^. Previous EEG–fMRI and source-imaging studies have associated microstate topographies with distributed functional networks, although these relationships are not necessarily exclusive ^33,34^. Consistent with previous neuromodulation studies reporting changes in microstate dynamics during pain modulation ^21,35,36^, we observed widespread training-related changes in microstate temporal characteristics and transition probabilities across feedback groups. Notably, the additional effect of authentic feedback was selective, with only the E→D transition showing a significant Training × Feedback authenticity interaction among the 20 directed transitions. Given previous associations of microstate E with salience-related processing and microstate D with attentional control ^30,31,33^, this reduction may reflect a decreased tendency for salience-related brain configurations to be followed by attention-control-related states. Such a shift could alter how salient information about impending nociceptive stimulation engages attentional resources during pain anticipation, although this functional interpretation requires further validation.

Importantly, greater training-related increases in somatosensory α power were associated with larger reductions in E→D transition probability, revealing a close relationship between oscillatory modulation and the temporal sequencing of preparatory brain states. Previous studies have demonstrated associations between α activity and EEG microstate characteristics ^42,43^, suggesting that local oscillatory activity and large-scale electrophysiological dynamics are closely interrelated. Within the gating-by-inhibition framework, increased somatosensory α activity may regulate cortical responsiveness to incoming sensory information ^10,44^. Our findings extend this perspective by suggesting that α enhancement may also be accompanied by changes in how preparatory brain states transition between configurations associated with salience processing and attentional control ^31,45^. Specifically, the association between α enhancement and reduced E→D transitions raises the possibility that stronger preparatory sensory regulation coincides with a reduced propensity for salience-related processing to engage subsequent attention-related states, potentially modifying the prioritization of impending nociceptive information. Although this relationship does not establish that α modulation directly drives microstate transitions, it suggests that feedback-dependent neural regulation may operate across complementary levels of brain organization, linking the modulation of local oscillatory activity with changes in the temporal dynamics of distributed neural states.

These changes in preparatory neural activity were accompanied by altered responses to subsequent nociceptive stimulation. Across feedback groups, active training was associated with reduced subjective ratings of pain intensity and unpleasantness, accompanied by attenuation of the N2 and P2 components of LEPs. These components are associated with salience detection and attentional orienting to nociceptive stimuli, with contributions from operculo-insular, anterior cingulate, and somatosensory cortices ^46,47^. Such general training-related effects are consistent with the established influence of attention, imagery, and other cognitive processes on pain perception ^48^. Importantly, N2 and P2 responses are not exclusively specific to nociceptive processing; they are also sensitive to stimulus salience and attentional orienting ^49,50^. Their attenuation may therefore reflect reduced cortical engagement with incoming nociceptive stimuli. Beyond these general training effects, authentic EEG-contingent feedback produced greater reductions in pain intensity and N2/P2 amplitudes than sham feedback, whereas no additional effect was observed for pain unpleasantness. These findings suggest that authentic feedback provides additional benefits for modulating subsequent pain-related cortical responses and subjective pain intensity beyond the general effects of active training. Nevertheless, despite the factorial design and measures taken to minimize nonspecific influences, expectancy-related effects cannot be entirely excluded, as participants’ expectations were not directly assessed.

The convergence of preparatory and pain-evoked neural changes raises an important question about how regulation of ongoing brain activity relates to subsequent pain processing. Our exploratory structural equation model identified significant indirect pathways linking feedback authenticity to changes in pain perception through somatosensory α modulation, E→D microstate transitions, and pain-evoked P2 responses. These associations suggest a potential cross-level pathway through which feedback-dependent regulation of preparatory brain states may influence the cortical processing of subsequent nociceptive events. However, changes in somatosensory α power were not directly associated with pain intensity reductions across participants, consistent with previous evidence that successful α regulation does not necessarily translate into analgesia ^19,20,51,52^. Thus, rather than supporting a simple correspondence between α enhancement and pain reduction, our findings point toward a more complex relationship involving preparatory oscillatory activity, brain state dynamics, and subsequent cortical responses. Although the proposed pathways remain to be causally established, they provide a mechanistically informed framework for understanding how closed-loop neurofeedback may bias subsequent pain processing.

The present findings extend current models of pain modulation by moving beyond the observation of pre-stimulus neural correlates toward the experimental regulation of preparatory brain states. Previous studies have demonstrated that spontaneous fluctuations in pre-stimulus neural activity predict subsequent pain responses ^3–5,7–9^, while neuromodulation studies have shown that externally manipulating somatosensory α oscillations can influence pain-evoked cortical responses and subjective pain ^8,36^. Building on this work, our findings demonstrate that preparatory neural activity can also be modulated through active, EEG-guided self-regulation, with authentic feedback producing additional changes in neural dynamics and subsequent pain processing. These results provide experimental support for a state-dependent account of pain, in which the configuration of ongoing brain activity contributes to how incoming nociceptive information is processed ^3,5,53^. Importantly, somatosensory α oscillations and EEG microstate dynamics capture complementary aspects of preparatory neural organization: somatosensory α oscillations index local inhibitory gain control ^10^, whereas EEG microstate dynamics capture how such local modulation is expressed within coordinated, large-scale brain organization ^35,36^. Their observed associations, together with changes in pain-evoked cortical responses and subjective perception, suggest a multilevel framework through which preparatory neural regulation may influence subsequent pain processing.

From a translational perspective, these findings highlight the potential of targeting preparatory brain states for pain regulation. The additional benefits of authentic EEG-contingent feedback underscore the value of reliable neural feedback for enhancing targeted regulation and subsequent pain-related outcomes. Moreover, the observation that neurofeedback delivered before nociceptive stimulation influenced subsequent pain responses suggests opportunities for anticipatory interventions targeting neural states before predictable painful events ^8,35^. The accompanying changes in EEG microstate dynamics further suggest that measures beyond local oscillatory power may inform future adaptive, state-sensitive neurofeedback protocols. Although the durability and clinical relevance of these effects remain to be established, our findings provide a foundation for developing preparatory brain-state-based approaches to pain modulation.

In summary, this study demonstrates that preparatory brain states can be experimentally modulated through closed-loop neurofeedback, with corresponding changes in subsequent pain processing and perception. Although active training was associated with broad neural and behavioral changes, authentic EEG-contingent feedback produced additional enhancement of somatosensory α activity, selective alterations in microstate transitions, and greater reductions in pain-evoked cortical responses and pain intensity. The observed associations among oscillatory activity, microstate dynamics, and pain responses further suggest that preparatory neural regulation involves multiple, interconnected levels of brain organization. These findings extend state-dependent accounts of pain from the prediction of sensory experience toward its experimental modulation, highlighting preparatory brain states as promising targets for future pain-regulation strategies.

## Methods

### Participants

An *a priori* sample size estimation was conducted using G*Power 3.1 ^54^ for a repeated-measures ANOVA with two groups (effect size = 0.25, α = 0.05, power = 0.95), indicating a minimum of 74 participants. To account for potential attrition, 80 healthy adults (42 males, 38 females; mean age ± SEM = 20.19 ± 0.25 years; Asian Chinese descent) were recruited through campus advertisements at a local university and randomly assigned to either the real-feedback group (*n* = 40) or the sham-feedback group (*n* = 40). Inclusion criteria required right-handedness and normal or corrected-to-normal vision. Exclusion criteria included: (1) acute or chronic pain, (2) cardiovascular or neurological disorders, (3) psychiatric diagnoses, (4) current use of psychoactive medication, or (5) any neuromodulation interventions within the past month. Data from six participants were excluded due to withdrawal from the study (*n* = 2) or excessive EEG artifacts (*n* = 4), resulting in a final analytic sample of 74 participants (real feedback: *n* = 38; sham feedback: *n* = 36). All participants provided written informed consent in accordance with the Declaration of Helsinki, and the study protocol was approved by the local ethics committee.

### General experimental procedure

This double-blind, sham-controlled study employed a 2 × 2 mixed factorial design, with autonomous training (training vs. nontraining) as a within-subject factor and feedback authenticity (real vs. sham) as a between-subject factor. This design enabled dissociation of the contributions of participants’ active engagement in self-regulation from the authenticity of the neurofeedback signal in shaping training-related pain modulation. To maintain blinding and minimize potential expectancy effects, participants were informed that they would perform a brain-control task and were instructed to use mental strategies to keep the feedback ball from falling. Neutral instructions were provided throughout the experiment, without disclosing the authenticity of the feedback or the hypothesized analgesic effects of neurofeedback. The study procedures were coordinated by two independent researchers with separate responsibilities. A blinded assessor conducted participant assessments, data collection, and data analysis, whereas an unblinded technician generated the random allocation sequence, assigned participants to the study groups, and delivered the real or sham neurofeedback intervention. The technician had no further contact with participants after the intervention. All other experimental procedures were identical across groups.

All participants completed the experimental procedures in a neuromodulation laboratory at a university in China. Upon arrival, participants completed a battery of psychometric questionnaires assessing individual differences in pain-related cognition, affect, and regulation. These included measures of pain catastrophizing (Pain Catastrophizing Scale), pain sensitivity (Pain Sensitivity Questionnaire), pain-related fear and anxiety (Fear of Pain Questionnaire; Pain Anxiety Symptoms Scale), perceived coping ability (Pain Resistance Scale), trait and state anxiety (State–Trait Anxiety Inventory), and emotion regulation strategies (Emotion Regulation Questionnaire). Following baseline assessment, the unblinded technician used a covariate-adaptive randomization procedure ^55^ to generate the allocation sequence and assign participants to the real- or sham-feedback groups, balancing demographic characteristics and psychometric profiles across conditions.

As illustrated in Fig. 1A, the experimental session began with a pain familiarization phase, during which participants received laser stimuli of varying intensities (2.5–4.5 J) and rated five 3.5 J stimuli using a 0–10 numerical rating scale (NRS) to confirm tolerability. Each subsequent condition (training and nontraining) comprised two components: (1) a 3-min resting-state EEG recording used to determine individualized neurofeedback thresholds; and (2) an NFB–pain rating task assessing the effects of feedback training on subsequent pain perception. For the real-feedback group, thresholds were derived from each participant’s own resting-state EEG data. For the sham-feedback group, thresholds were generated from a resting-state EEG database consisting of recordings from three independent individuals not involved in the study. Each database participant contributed five 5-min resting blocks, from which 3-min segments were randomly selected to define thresholds separately for the nontraining and training conditions.

As shown in Fig. 1B, each NFB–pain rating trial began with a 16-s neurofeedback phase, during which the position of a visual feedback stimulus (a moving ball) was updated every 400 ms based on real-time EEG processing. In the nontraining condition, participants passively observed the feedback display, whereas in the training condition, they actively attempted to regulate their brain activity to prevent the ball from falling. Following a fixation interval (3–5 s), a noxious laser stimulus (3.5 J; wavelength = 1.34 μm; pulse duration = 4 ms; beam diameter = 7 mm; Electronical Engineering, Italy) was delivered to the dorsal web space of the left hand. After a further delay (3–5 s), participants verbally rated pain intensity and unpleasantness on separate 0–10 NRS scales (0 = none, 10 = maximum). Each trial ended with a 2–4 s intertrial interval. Participants completed 30 trials per condition (training and nontraining), divided into two blocks, yielding a total task duration of approximately 25 min per condition.

### Neurofeedback training protocol

The NFB system provided continuous visual feedback by mapping somatosensory α-power onto the horizontal position of a white ball displayed on a gray background (Fig. 1C). The closed-loop architecture comprised four components: EEG acquisition, real-time signal processing, visual feedback, and participant self-regulation ^19,20,56,57^.

**(1) *EEG acquisition*** EEG data were recorded using a 64-channel Ag/AgCl electrode cap (Brain Products GmbH, Germany) arranged according to the international 10–20 system. Signals were referenced online to CPz, sampled at 1000 Hz, and bandpass filtered online between 0.01 and 100 Hz. Electrode impedances were kept below 10 kΩ. Control signals were derived from electrodes over the right somatosensory cortex (C4, CP4, CP6), contralateral to the site of pain stimulation.
**(2) *Real-time processing.*** Every 400 ms, an 800-ms segment of the most recent EEG data (800 samples at 1000 Hz) was extracted from a continuous data buffer. For each of the three target electrodes (C4, CP4, and CP6), the signal was mean-centered (constant detrending) and band-pass filtered between 1 and 40 Hz using a third-order Butterworth filter with forward–reverse filtering (*filtfilt*). The filtered signal was then transformed using an 800-point fast Fourier transform (FFT), yielding a frequency-bin spacing of 1.25 Hz. Alpha-band power (8–13 Hz) was calculated by averaging power values across the FFT frequency bins falling within this range for each electrode. The resulting α-power values were then averaged across C4, CP4, and CP6 to obtain the real-time α-band feedback index.
**(3) *Visual feedback***. In the real-feedback group, an individualized threshold was determined from the mean absolute α power during a 3-min resting-state EEG recording obtained immediately before each condition. During neurofeedback, the real-time α-power index was compared with this threshold every 400 ms to control ball movement. When α power exceeded the threshold, the ball moved one step to the right; otherwise, it moved one step diagonally downward and to the right. In the sham-feedback group, visual feedback was generated using a randomly selected 16-s EEG segment from a resting-state EEG database of independent donors for each trial. Consequently, sham feedback was independent of participants’ ongoing EEG activity.
**(4) *Self-regulation***. In the training condition, participants were instructed to “mentally keep the ball from falling,” without explicit strategy guidance, encouraging spontaneous self-regulation. In the nontraining condition, participants passively observed the ball’s movement without attempting regulation.

This closed-loop system enabled real-time translation of somatosensory α-power into dynamic visual feedback under tightly controlled conditions. Following the experiment, a blinding check assessed perceived feedback authenticity by asking participants whether they believed they had controlled the ball’s movement during training.

### Offline EEG data processing

Offline EEG preprocessing was performed using EEGLAB ^58^. Two EEG datasets were extracted for separate analyses: (1) neurofeedback-period EEG recorded during the 16-s feedback phase of each training and nontraining trial, preceding the fixation interval and pain stimulation; and (2) pain-evoked EEG recorded around the onset of the laser stimulus. The two datasets were processed separately because they required different epoch definitions, but the same core preprocessing and artifact-correction procedures were applied.

For both datasets, EEG signals were band-pass filtered between 1 and 100 Hz and notch filtered at 50 Hz to remove line noise. Neurofeedback-phase EEG was segmented into consecutive, non-overlapping 2-s epochs within each 16-s feedback interval, whereas pain-evoked EEG was segmented into 1500-ms epochs time-locked to laser onset (−500 to 1000 ms). Within each dataset, training and nontraining epochs were pooled for preprocessing. Epochs containing prominent movement-related or electrode artifacts were visually rejected, followed by independent component analysis (ICA) ^59^ to remove ocular, muscle, and other non-neural artifacts. The corrected EEG data were then re-referenced to the common average, with the original reference electrode (CPz) restored to the montage.

### EEG oscillatory power analysis

To quantify training-related changes in EEG oscillatory activity, power spectral density (PSD) was estimated from non-overlapping 2-s epochs during the neurofeedback phase using the FieldTrip toolbox ^60^. PSDs were computed using a multitaper fast Fourier transform (FFT) with discrete prolate spheroidal sequence (DPSS) tapers, yielding 1-Hz spectral smoothing over the 1–100 Hz frequency range (0.5-Hz frequency resolution). Power values were log-transformed to decibel (dB) units and averaged across epochs within each condition for each participant.

Band-limited power was extracted for five canonical frequency bands ^1,35,61^: delta (δ, 1–4 Hz), theta (θ, 4–7 Hz), alpha (α, 8–13 Hz), beta (β, 13–30 Hz), and gamma (γ, 30–90 Hz). For regional analyses, electrodes were grouped into four predefined regions of interest (ROIs): target somatosensory (C4, CP4, CP6), nontarget somatosensory (C3, CP3, CP5), frontal (F1, Fz, F2), and parietal (P1, Pz, P2) regions. Band-limited power was averaged across electrodes within each ROI, separately for the training and nontraining conditions.

To characterize the temporal dynamics of α modulation, target-S1 α power was additionally examined during the first 8 s, last 8 s, and final 2 s of the 16-s neurofeedback phase. To assess whether feedback-dependent α modulation persisted immediately before nociceptive stimulation, α power was also quantified during the final 2 s preceding laser onset, within the fixation interval. The same spectral estimation procedures were applied to these time windows, and α power was analyzed separately for each participant and condition.

### Source-space α-power and functional connectivity analyses

Source reconstruction was performed on the preprocessed EEG data recorded during the neurofeedback phase using a linearly constrained minimum variance (LCMV) beamformer ^62^. A standard template boundary element method (BEM) head model was used to construct a 10-mm source grid. Electrode locations were obtained from the channel-coordinate information in each participant’s EEG dataset and transformed into the coordinate system of the source model using a consistent procedure across participants. Lead fields were computed with a rank-reduction setting of 3 and normalized. For each participant, a common spatial filter was constructed using α-band (8–13 Hz) data pooled across the training and nontraining conditions, with 5% regularization and fixed source orientation. The resulting common filter was then applied separately to the training and nontraining data to reconstruct condition-specific source-level EEG signals. This common-filter approach ensured that source reconstruction was performed using the same spatial filters across conditions, allowing subsequent comparisons of α-power and functional connectivity without introducing condition-specific differences in spatial filtering.

#### Source-space α-power analysis

Spectral power was estimated separately for the training and nontraining conditions within the α-frequency band (8–13 Hz) using FieldTrip’s *mtmfft* method with DPSS tapers and a frequency smoothing of 2.5 Hz ^60^. Power estimates were averaged across trials. Source-level power was then mapped to the left and right postcentral gyri (PoCGs) defined by the Automated Anatomical Labeling (AAL) atlas ^63^. For each hemisphere, power values were averaged across all source points within the corresponding PoCG and then log-transformed to decibel (dB) units. This yielded one α-power value for each participant, hemisphere, and condition.

#### Source-space functional connectivity analysis

Interhemispheric α-band functional connectivity was quantified using the dwPLI ^64,65^ between the right and left PoCGs, corresponding to the target and nontarget S1, respectively. Equal numbers of artifact-free epochs were randomly selected from the training and nontraining conditions for each participant. Source-reconstructed signals were decomposed using a Fourier transform with a Hanning taper, retaining trial-level Fourier coefficients. The dwPLI was calculated across 8–13 Hz using FieldTrip’s *wpli_debiased* method ^60^. Frequency-specific dwPLI values were averaged across the α band, followed by averaging across all source-point pairs connecting the right and left PoCGs. This yielded one interhemispheric α-band connectivity value for each participant and condition.

### EEG microstate analysis

EEG microstate analysis was conducted to assess how NFB influenced large-scale brain state dynamics, following established procedures ^35,36,66,67^. Microstate analysis was performed offline using EEG recorded during the 16-s neurofeedback phase across all training and nontraining trials. Each feedback period was divided into eight consecutive 2-s epochs. The preprocessed EEG signals were downsampled to 250 Hz and bandpass filtered between 2 and 20 Hz. Global field power (GFP) was computed as the spatial standard deviation across electrodes at each time point, and scalp topographies at GFP peaks were extracted to capture periods of maximal global activity and spatial stability.

Microstate maps were first identified separately for each participant and condition using GFP-peak-based atomize-and-agglomerate hierarchical clustering (AAHC). Within each feedback group, these individual maps were hierarchically aggregated to derive condition-level maps and subsequently a group-level template comprising five microstate classes (A–E). The real- and sham-feedback templates were constructed independently, without pooling data across groups. Despite this independent estimation, corresponding microstate classes showed high spatial similarity between groups (all *r* > 0.95, *p* < 0.001; Supplementary Fig. S2). Finally, each group-specific template was back-fitted to the training and nontraining EEG data of participants in the corresponding group to quantify microstate temporal characteristics and transition probabilities.

For each microstate class, three temporal parameters were computed ^30^: (1) the average duration that each microstate remains stable (i.e., mean duration; ms), (2) the frequency of occurrence for each microstate per second (i.e., occurrence; events/s), (3) the percentage of the total recording time occupied by each microstate (i.e., time coverage; %). In addition, transition probabilities were calculated to quantify the likelihood of switching between microstate classes, normalized by the total number of transitions. Transition matrices were computed separately for each participant and condition using the group-level microstate templates.

### Laser-evoked potential (LEP) analysis

To examine cortical responses to nociceptive stimulation, the preprocessed pain-locked EEG was additionally low-pass filtered at 30 Hz and baseline-corrected using the −500 to 0 ms pre-stimulus interval. For each participant, laser-evoked EEG responses were averaged separately for the training and nontraining conditions, yielding two condition-specific waveforms. Group-level grand averages were subsequently computed to visualize representative LEP waveforms and spline-interpolated scalp topographies.

Analyses focused on the N2 and P2 components ^8,36^, which exhibited maximal amplitudes over central electrodes (C1, Cz, C2, FC1, FCz, FC2, CP1, CPz, and CP2). The N2 component was defined as the most negative deflection within 180–300 ms following stimulus onset, and the P2 component as the most positive deflection within 300–500 ms. For each participant and condition, peak amplitudes of N2 and P2 were extracted within these predefined latency windows.

### Statistical analysis

Statistical analyses were performed on 74 participants (real feedback: *n* = 38; sham feedback: *n* = 36) using MATLAB (MathWorks, Natick, MA, USA), IBM SPSS Statistics (Version 19; IBM Corp., Armonk, NY, USA), and IBM SPSS Amos (Version 26; IBM Corp., Armonk, NY, USA). All statistical tests were two-sided where applicable, with a significance threshold of *p* < 0.05. Baseline differences between the real- and sham-feedback groups were assessed using χ² tests for categorical variables and independent-samples *t*-tests for continuous measures.

To evaluate training-related effects and their dependence on feedback authenticity, two-way mixed-design ANOVAs were conducted with Training (training vs. nontraining) as a within-subject factor and Feedback authenticity (real vs. sham) as a between-subject factor. These analyses were applied to sensor-level oscillatory power, source-level α power and functional connectivity, EEG microstate characteristics (mean duration, occurrence, time coverage, and transition probabilities), pain ratings, and LEP amplitudes (N2 and P2). The main effect of Training assessed the overall difference between training and nontraining conditions across feedback groups, whereas the Training × Feedback authenticity interaction tested whether this training-related difference varied between real and sham feedback. Importantly, the interaction provided the primary test of additional effects associated with authentic EEG-contingent feedback beyond those observed with sham feedback, which may reflect general effects of active engagement, attention, and other nonspecific factors. Significant interactions were followed by simple-effects comparisons. Training-related change scores (Δ = Training − Nontraining) were calculated for each participant and compared between feedback groups. To account for multiple comparisons, FDR ^37^ using the Benjamini–Hochberg procedure was applied separately to EEG oscillatory power, microstate temporal characteristics, and transition probabilities. FDR-adjusted *p* values are reported as *p_FDR_*.

Additional analyses examined the temporal dynamics and robustness of the main findings. Target-S1 α power was assessed across different time windows during the neurofeedback and pre-stimulus periods, while within-session changes were examined using three-way mixed-design ANOVAs incorporating Training, Phase (early vs. late trials), and Feedback authenticity. Associations between neural modulation and pain perception were evaluated using Pearson correlations of participant-level training-related changes and trial-level linear mixed-effects models (LMMs). For trial-level analyses, α power was centered within each participant and condition, with α power, Training, Feedback authenticity, and their interactions included as fixed effects and participant-specific random intercepts. Additional multiband models incorporated power across five frequency bands to assess the specificity of α–pain associations.

Finally, to integrate neural and behavioral changes across successive stages of pain processing, we conducted an exploratory SEM analysis. Whereas the preceding analyses examined individual associations between neural and behavioral measures, the SEM provided an integrative framework for examining the hypothesized relationships among preparatory neural modulation, subsequent pain-evoked cortical responses, and subjective pain perception. Feedback authenticity (real vs. sham) was specified as the exogenous variable, while participant-level training-related changes (Training − Nontraining) in somatosensory α power, E→D microstate transition probability, LEP P2 amplitude, and pain ratings were included as observed variables. The model was designed to examine both direct and indirect associations linking feedback authenticity to pain-related outcomes through preparatory neural changes. Model fit was evaluated using the χ² statistic, RMSEA, RMR, and GFI. Indirect effects were assessed using bootstrapped 95% confidence intervals, with statistical significance determined by intervals excluding zero.

## Supporting information

CRED-nf Checklist

## Acknowledgements

This work was supported by the ‘Sci-Tech Innovation 2030’ Brain Science and Brain-Inspired Intelligence Technology Research by the Ministry of Science and Technology of China (grant number 2022ZD0206400), National Natural Science Foundation of China (grant numbers 62276262 and 32671449), Shenzhen Basic Research Project (grant numbers QNXMB20250701094310014 and JCYJ20230808105805012), and Shenzhen University 2035 Program for Excellent Research (grant number 2024C003).

## Data and code availability

The raw EEG data, the processed data that underlie the statistical analyses and figures, and the custom MATLAB scripts used for preprocessing and analysis are publicly available in the Open Science Framework repository (https://osf.io/x4gf2) ^68^.

## Authorship contributions

YH.Z.: Formal analysis, Visualization; Writing-original draft, Writing-review and editing. S.Q.: Conceptualization, Formal analysis, Writing-review and editing. QQ.Z.: Conceptualization, Data curation, Formal analysis. SH.C.: Formal analysis, Writing-review and editing. J.L.: Formal analysis, Writing-review and editing. XY. L.: Conceptualization, Writing-review and editing. RC.J.: Conceptualization, Writing-review and editing. WW.P.: Conceptualization; Formal analysis, Writing-original draft, Writing-review and editing, Funding acquisition, Supervision. All authors read and approved the final manuscript.

## Competing interests

The authors declare no competing interests.

## Supplementary information

## Section 1. Participant-reported neurofeedback regulation strategies

**Table S1.** Self-reported regulation strategies during neurofeedback training.

| Strategy category | Coding criteria | Representative examples | *n* (%) |
|---|---|---|---|
| **Memory and imagery** | Recalling past experiences or imagining scenes and events. | Recalling emotional experiences; imagining social interactions, sports, or everyday activities. | 64 (86.49) |
| **Attentional focus** | Directing attention to external feedback or ongoing internal sensations without deliberately modifying them. | Focusing on ball movement; attending to breathing, heartbeat, or bodily sensations. | 39 (52.70) |
| **Somatic or physiological regulation** | Deliberately modifying breathing, muscle activity, or bodily movements. | Adjusting breathing; changing muscle tension; moving limbs or controlling eye movements. | 35 (47.30) |
| **Verbal or numerical cognitive strategies** | Engaging in internally generated verbal or numerical mental operations. | Counting; mental arithmetic; silent recitation; repeating words or phrases. | 34 (45.95) |
| **Other strategies** | Self-reported approaches not clearly classified into the above categories. | Thought suppression; treating the task as a game; other individual approaches. | 22 (29.73) |

**Note.** Strategies were categorized from participants' self-reports collected after neurofeedback training. Participants were not instructed to use specific strategies. Categories were non-mutually exclusive, but each participant was counted only once per category. Frequencies and percentages are based on the 74 participants who provided strategy reports and therefore do not sum to 100%.

## Section 2. Supplementary analyses of EEG oscillatory activity

**Table S2.**
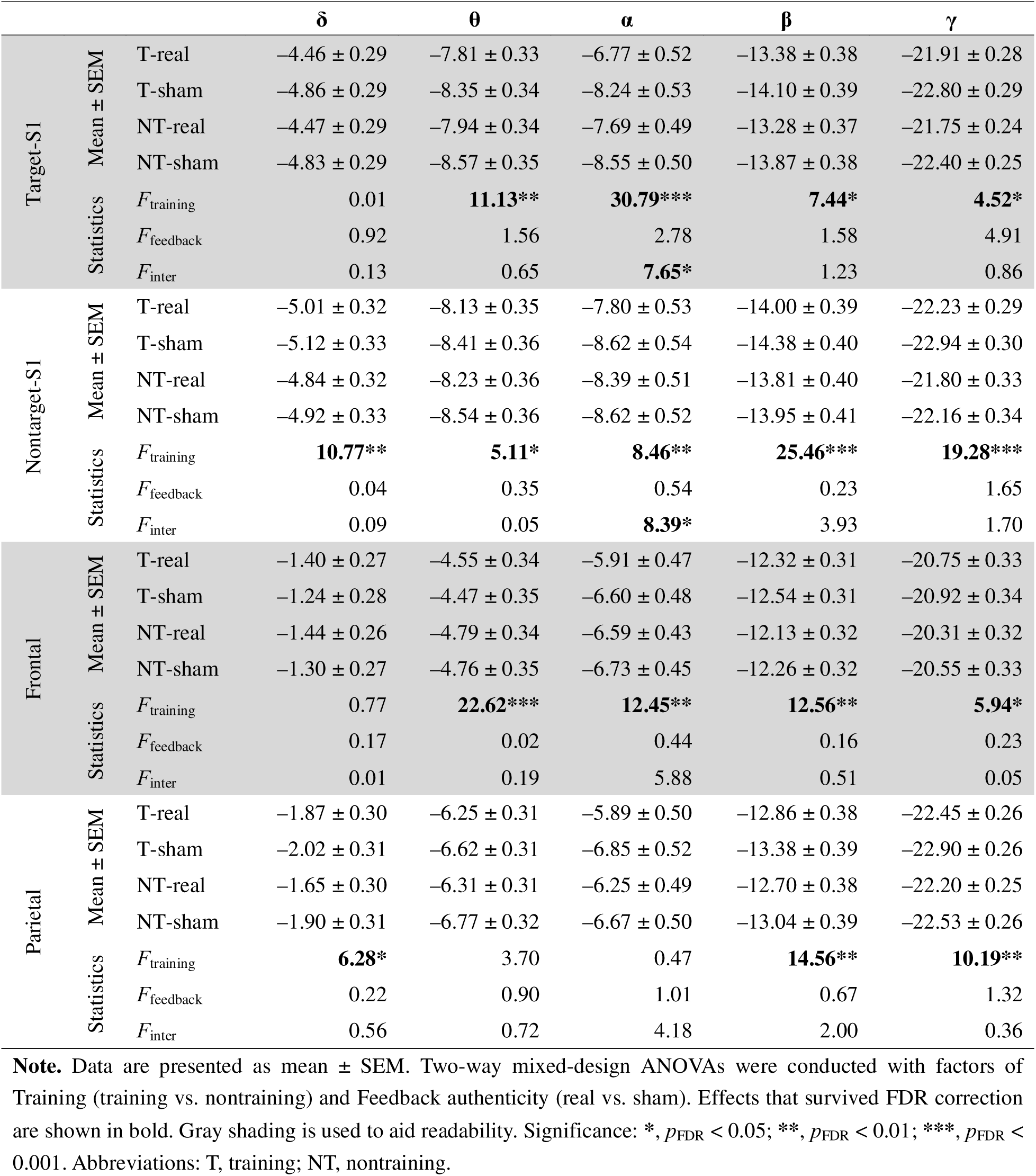
Statistics for brain oscillation power during the NFB phase.

**Figure S1.**
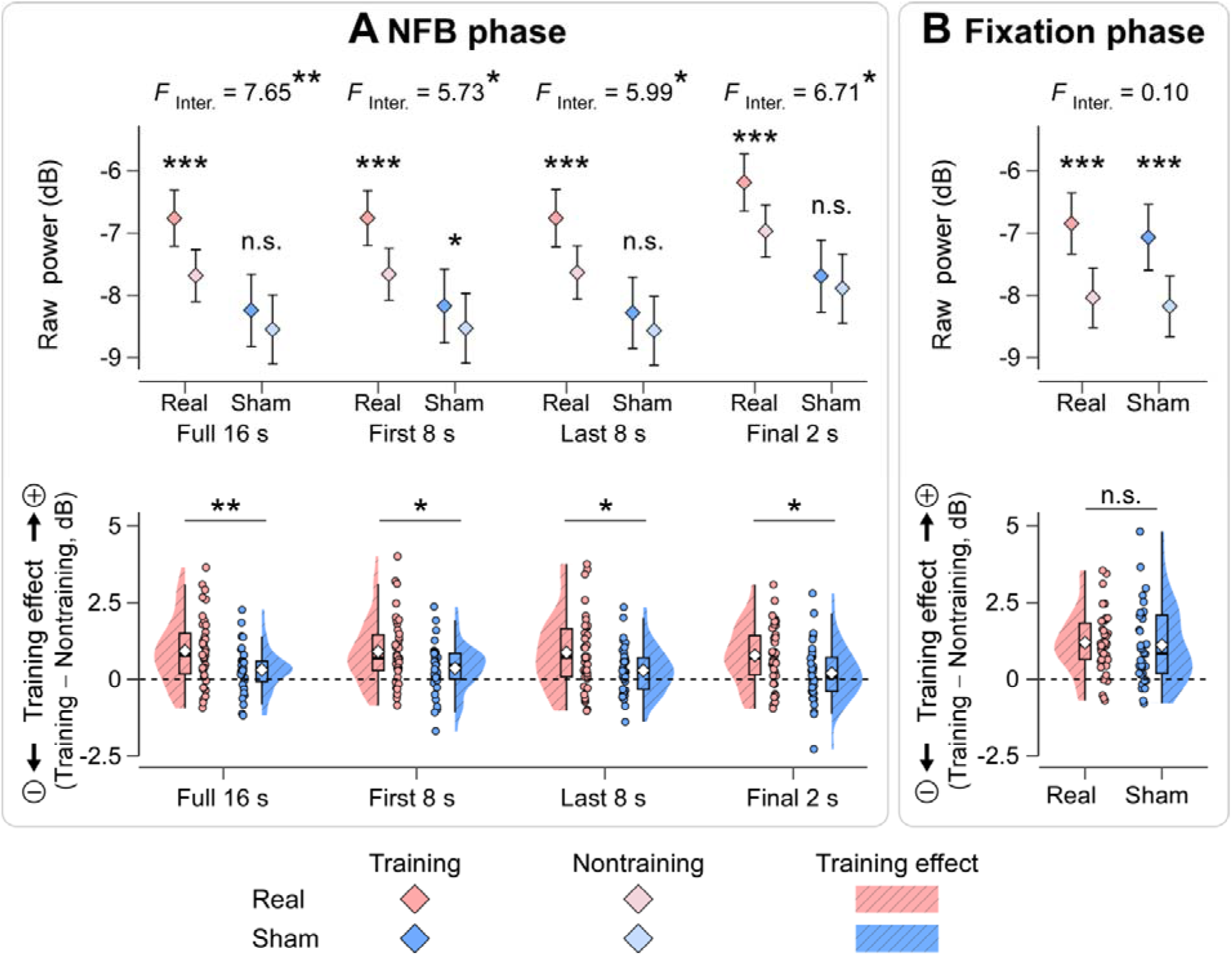
Temporal profile of target-S1 α modulation during neurofeedback and before pain delivery. ***(A) Neurofeedback phase.*** Target-S1 α power during training and nontraining is shown for the real- and sham-feedback groups across four windows: full 16 s, first 8 s, last 8 s, and final 2 s. The corresponding training-related changes (Training − Nontraining) show significant Training × Feedback authenticity interactions across all four windows. ***(B) Pre-pain fixation phase.*** Target-S1 α power and the corresponding training-related changes are shown for the final 2 s immediately before laser stimulation. Although α power increased during training in both groups, the Training × Feedback authenticity interaction was not significant. Real and sham feedback are shown in red and blue, respectively. In the upper panels, diamonds indicate means. In the lower panels, violin plots show data distributions; box plots indicate medians and interquartile ranges; diamonds indicate means; dots represent individual participants. Significance: **\***, *p* < 0.05; **\*\***, *p* < 0.01; **\*\*\***, *p* < 0.001; n.s., not significant. Abbreviations: Inter., Training × Feedback authenticity interactions; “+” indicates a greater training effect, whereas “−” indicates a smaller training effect.

## Section 3. Supplementary analyses of EEG microstate dynamics

**Figure S2.**
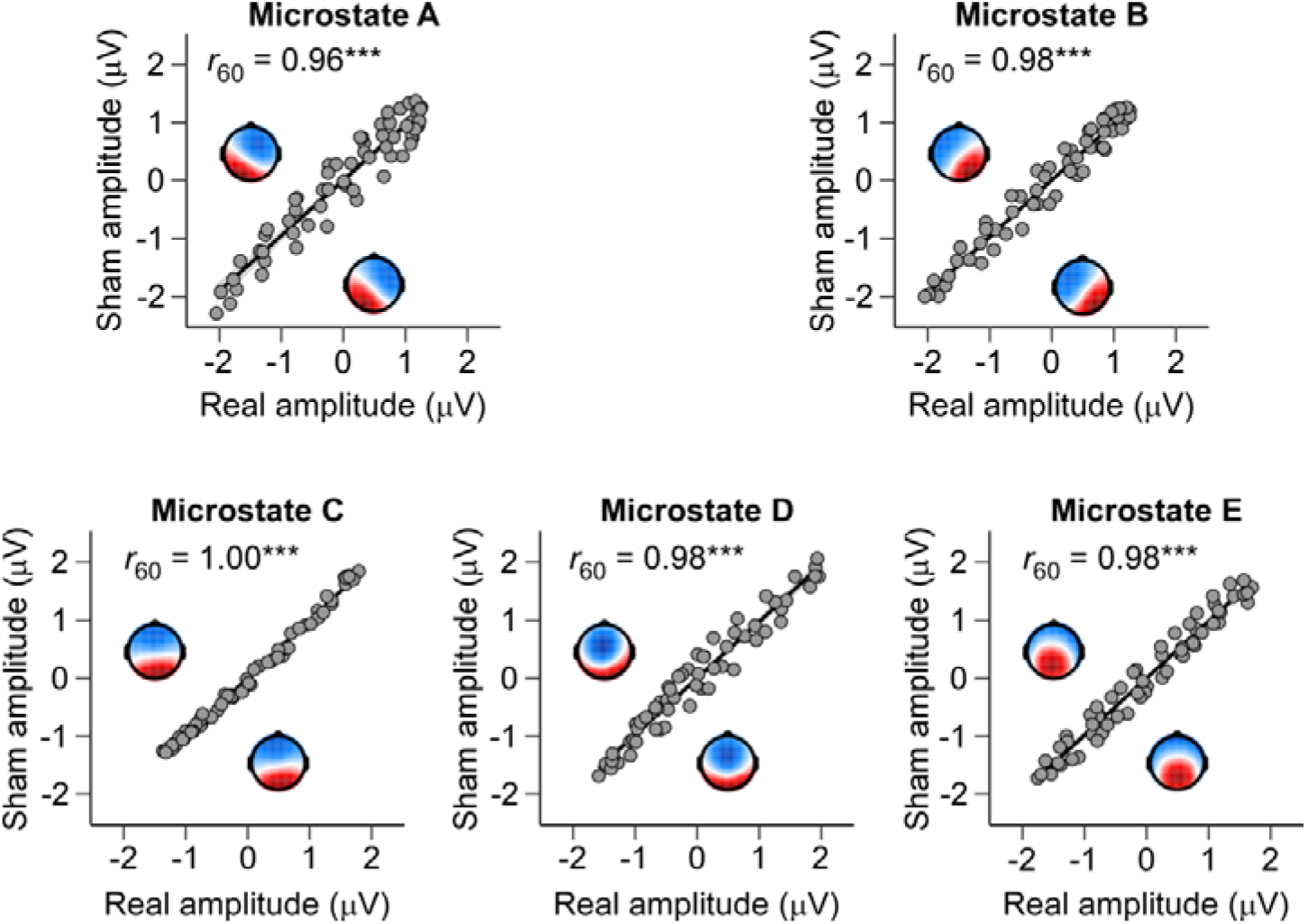
Spatial correspondence of EEG microstate topographies between real- and sham-feedback groups. Group-level scalp maps and corresponding electrode-wise spatial correlations are shown for microstates A–E. Scatterplots compare normalized scalp potentials between the real- and sham-feedback groups, with each point representing one electrode. Black lines indicate linear regression fits, with shaded areas representing 95% confidence intervals. All five microstate classes showed high spatial similarity between groups (all *r*_60_ > 0.95, *p* < 0.001), supporting the correspondence of microstate classes derived independently in the two feedback groups. Significance levels: **\*\*\***, *p* < 0.001.

**Figure S3.**
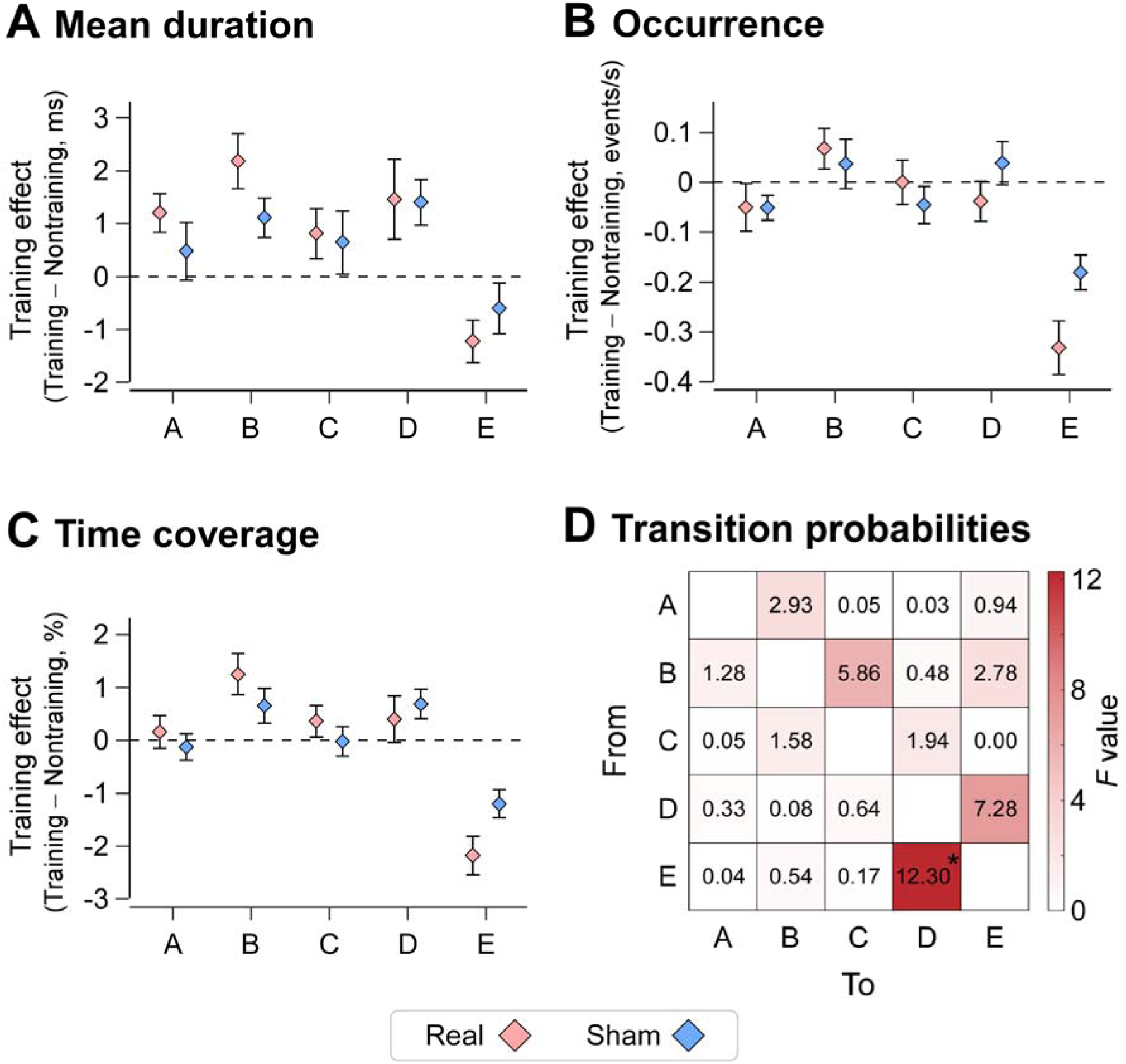
Training-related changes in EEG microstate temporal characteristics and transition probabilities. *(A–C) Microstate temporal characteristics.* Training-related changes (Training − Nontraining) in (A) mean duration, (B) occurrence, and (C) time coverage are shown for microstates A–E separately in the real- and sham-feedback groups. No Training × Feedback authenticity interaction remained significant after FDR correction across these measures. Red and blue diamonds represent the means for the real- and sham-feedback groups, respectively, and error bars indicate standard errors of the mean (SEM). *(D) Microstate transition probabilities.* The matrix shows the *F* statistics for the Training × Feedback authenticity interaction across all 20 directed transitions among microstates A–E. Each cell represents the *F* statistic from the corresponding 2 × 2 mixed-design ANOVA, with rows indicating the originating microstates and columns indicating the destination microstates. Only the E→D transition showed a significant interaction after FDR correction. Rows indicate the originating microstate (From), and columns indicate the destination microstate (To). For example, the cell in the first row and second column represents the A→B transition, whereas the cell in the fourth row and third column represents the D→C transition. Diagonal cells representing self-transitions were excluded. Color intensity represents the magnitude of the *F* statistic, with darker red indicating larger *F* values. Significance levels: *, *p_FDR_*< 0.05.

**Table S3.** Statistics for EEG microstate temporal characteristics during the NFB phase.

|  |  | **Training condition** |  | **Nontraining condition** |  | **Statistics** |  |  |
|---|---|---|---|---|---|---|---|---|
|  |  | Real | Sham | Real | Sham | *F*<sub>Training</sub> | *F*<sub>Feedback</sub> | *F*<sub>Inter.</sub> |
| Mean duration | Microstate A | 60.71 ± 1.10 | 59.54 ± 1.13 | 59.51 ± 1.15 | 59.06 ± 1.18 | **6.70\*** | 0.27 | 1.24 |
|  | Microstate B | 59.58 ± 1.06 | 58.41 ± 1.09 | 57.40 ± 0.93 | 57.29 ± 0.96 | **25.95\*\*\*** | 0.22 | 2.75 |
|  | Microstate C | 56.96 ± 1.30 | 57.69 ± 1.33 | 56.14 ± 1.16 | 57.05 ± 1.20 | 3.77 | 0.23 | 0.05 |
|  | Microstate D | 57.80 ± 1.73 | 56.42 ± 1.78 | 56.34 ± 1.44 | 55.02 ± 1.48 | **10.48\*\*** | 0.38 | 0.004 |
|  | Microstate E | 56.74 ± 1.21 | 57.76 ± 1.25 | 57.97 ± 1.23 | 58.36 ± 1.26 | **8.54\*** | 0.17 | 1.01 |
| Occurrence | Microstate A | 3.67 ± 0.12 | 3.48 ± 0.12 | 3.72 ± 0.11 | 3.53 ± 0.12 | 3.47 | 1.42 | < 0.001 |
|  | Microstate B | 3.63 ± 0.12 | 3.62 ± 0.13 | 3.56 ± 0.12 | 3.58 ± 0.12 | 2.69 | 0.003 | 0.23 |
|  | Microstate C | 2.69 ± 0.11 | 2.78 ± 0.11 | 2.69 ± 0.11 | 2.82 ± 0.12 | 0.60 | 0.51 | 0.60 |
|  | Microstate D | 3.32 ± 0.12 | 3.47 ± 0.12 | 3.36 ± 0.11 | 3.43 ± 0.12 | < 0.001 | 0.42 | 1.70 |
|  | Microstate E | 3.65 ± 0.13 | 3.91 ± 0.14 | 3.98 ± 0.12 | 4.09 ± 0.13 | **62.08\*\*\*** | 1.02 | 5.41 |
| Time coverage | Microstate A | 22.26 ± 0.95 | 20.74 ± 0.98 | 22.10 ± 0.92 | 20.87 ± 0.94 | 0.01 | 1.11 | 0.52 |
|  | Microstate B | 21.75 ± 0.95 | 21.08 ± 0.98 | 20.50 ± 0.84 | 20.42 ± 0.86 | **14.01\*\*** | 0.09 | 1.35 |
|  | Microstate C | 15.61 ± 1.00 | 16.15 ± 1.03 | 15.25 ± 0.95 | 16.17 ± 0.98 | 0.69 | 0.29 | 0.89 |
|  | Microstate D | 19.58 ± 1.16 | 19.52 ± 1.19 | 19.18 ± 1.02 | 18.83 ± 1.05 | 4.22 | 0.02 | 0.30 |
|  | Microstate E | 20.80 ± 1.06 | 22.52 ± 1.09 | 22.98 ± 1.00 | 23.71 ± 1.03 | **54.61\*\*\*** | 0.73 | 4.59 |

**Note.** Data are presented as mean ± SEM. Two-way mixed-design ANOVAs were conducted with factors of Training (training vs. nontraining) and Feedback authenticity (real vs. sham). Effects that survived FDR correction are shown in bold. Significance: \*, *p*<sub>FDR</sub> < 0.05; \*\*, *p*<sub>FDR</sub> < 0.01; \*\*\*, *p*<sub>FDR</sub> < 0.001. Abbreviations: Inter., Training × Feedback authenticity interactions.

**Table S4.** Statistics for EEG microstate transition probabilities during the NFB phase.

|  | Training condition |  | Nontraining condition |  | Statistics |  |  |
|---|---|---|---|---|---|---|---|
|  | Real | Sham | Real | Sham | *F*<sub>Training</sub> | *F*<sub>Feedback</sub> | *F*<sub>Inter.</sub> |
| A → B | 0.062 ± 0.003 | 0.053 ± 0.004 | 0.058 ± 0.003 | 0.052 ± 0.003 | **7.05\*** | 2.76 | 2.93 |
| B → A | 0.062 ± 0.004 | 0.054 ± 0.004 | 0.058 ± 0.003 | 0.052 ± 0.003 | **6.74\*** | 2.24 | 1.28 |
| A → C | 0.037 ± 0.002 | 0.037 ± 0.002 | 0.037 ± 0.002 | 0.037 ± 0.002 | 0.005 | 0.003 | 0.05 |
| C → A | 0.037 ± 0.002 | 0.037 ± 0.002 | 0.036 ± 0.002 | 0.037 ± 0.002 | 0.07 | < 0.001 | 0.05 |
| A → D | 0.050 ± 0.002 | 0.046 ± 0.002 | 0.049 ± 0.002 | 0.045 ± 0.002 | 0.24 | 1.58 | 0.03 |
| D → A | 0.050 ± 0.002 | 0.046 ± 0.002 | 0.050 ± 0.002 | 0.045 ± 0.002 | 1.54 | 2.21 | 0.33 |
| A → E | 0.059 ± 0.004 | 0.059 ± 0.004 | 0.063 ± 0.003 | 0.061 ± 0.003 | **10.34\*\*** | 0.08 | 0.94 |
| E → A | 0.059 ± 0.004 | 0.058 ± 0.004 | 0.063 ± 0.003 | 0.062 ± 0.003 | **14.03\*\*** | 0.03 | 0.04 |
| B → C | 0.038 ± 0.002 | 0.037 ± 0.002 | 0.035 ± 0.002 | 0.037 ± 0.002 | **7.22\*** | 0.07 | 5.86 |
| C → B | 0.038 ± 0.002 | 0.038 ± 0.002 | 0.036 ± 0.002 | 0.037 ± 0.002 | **7.05\*** | 0.15 | 1.58 |
| B → D | 0.051 ± 0.003 | 0.051 ± 0.003 | 0.047 ± 0.003 | 0.047 ± 0.003 | **20.92\*\*\*** | 0.002 | 0.48 |
| D → B | 0.050 ± 0.003 | 0.051 ± 0.003 | 0.046 ± 0.002 | 0.048 ± 0.003 | **14.03\*\*** | 0.05 | 0.08 |
| B → E | 0.055 ± 0.003 | 0.060 ± 0.003 | 0.058 ± 0.003 | 0.061 ± 0.003 | **5.64\*** | 1.26 | 2.78 |
| E → B | 0.055 ± 0.003 | 0.060 ± 0.003 | 0.058 ± 0.003 | 0.061 ± 0.003 | **4.50\*** | 1.00 | 0.54 |
| C → D | 0.038 ± 0.003 | 0.039 ± 0.003 | 0.034 ± 0.002 | 0.037 ± 0.002 | **8.64\*\*** | 0.26 | 1.94 |
| D → C | 0.038 ± 0.003 | 0.040 ± 0.003 | 0.034 ± 0.002 | 0.038 ± 0.002 | **9.77\*\*** | 0.55 | 0.64 |
| C → E | 0.041 ± 0.003 | 0.043 ± 0.004 | 0.045 ± 0.004 | 0.046 ± 0.004 | **19.47\*\*\*** | 0.05 | < 0.001 |
| E → C | 0.041 ± 0.003 | 0.042 ± 0.003 | 0.044 ± 0.004 | 0.044 ± 0.004 | **13.01\*\*** | 0.01 | 0.17 |
| D → E | 0.051 ± 0.003 | 0.057 ± 0.003 | 0.057 ± 0.003 | 0.059 ± 0.003 | **26.08\*\*\*** | 1.04 | 7.28 |
| E → D | 0.051 ± 0.003 | 0.059 ± 0.003 | 0.057 ± 0.003 | 0.060 ± 0.003 | **26.44\*\*\*** | 1.62 | **12.30\*** |

**Note.** Data are presented as mean ± SEM. Two-way mixed-design ANOVAs were conducted with factors of Training (training vs. nontraining) and Feedback authenticity (real vs. sham). Effects that survived FDR correction are shown in bold. Significance: \*, *p*<sub>FDR</sub> < 0.05; \*\*, *p*<sub>FDR</sub> < 0.01; \*\*\*, *p*<sub>FDR</sub> < 0.001. Abbreviations: Inter., Training × Feedback authenticity interactions; "→" denotes the transition probability from one microstate to another.

## Section 4. Within-session temporal effects and robustness of feedback-dependent findings

**Table S5.** Within-session temporal effects on pain ratings and EEG responses.

|  | Phase | Training × Phase | Feedback × Phase | Training × Feedback | Training × Feedback × Phase |
|---|---|---|---|---|---|
| Pain intensity | **32.48\*\*\*** | **11.50\*\*** | 1.09 | **5.65\*** | 0.04 |
| Unpleasantness | **29.55\*\*\*** | **5.98\*** | 0.69 | 1.61 | 0.05 |
| N2 amplitude | **106.66\*\*\*** | **32.80\*\*\*** | 2.19 | **7.71\*\*** | < 0.001 |
| P2 amplitude | **60.36\*\*\*** | **22.36\*\*\*** | **4.20\*** | **7.24\*\*** | 0.81 |
| Target S1 α-power | **45.78\*\*\*** | **7.55\*\*** | 0.98 | **7.66\*\*** | < 0.001 |
| Nontarget S1 α-power | **29.29\*\*\*** | 1.13 | 0.87 | **8.40\*\*** | 0.14 |

**Note.** Each condition comprised 30 trials, divided into the first (Trials 1–15) and second (Trials 16–30) halves. A 2 (Training) × 2 (Phase) × 2 (Feedback authenticity) mixed-design ANOVA was conducted for each outcome, with Training and Phase as within-subject factors and Feedback authenticity as a between-subject factor. Values indicate *F* statistics. Training and Feedback authenticity main effects are omitted from this table to emphasize within-session temporal effects and the critical interactions. Significance: \*, *p* < 0.05; \*\*, *p* < 0.01; \*\*\*, *p* < 0.001.

## Section 5. Neural–behavioral associations across participant and trial levels

**Table S6.** Across-participant correlations between training-related changes in neurofeedback-period oscillatory power and pain intensity.

| **Frequency band** | **Target S1** | **Nontarget S1** | **Frontal** | **Parietal** |
|---|---|---|---|---|
| δ | 0.03 (0.80) | 0.16 (0.18) | 0.13 (0.27) | 0.05 (0.70) |
| θ | 0.26 (0.02) | 0.29 (0.01) | 0.27 (0.02) | 0.20 (0.09) |
| α | −0.09 (0.44) | 0.02 (0.83) | 0.01 (0.93) | 0.05 (0.67) |
| β | −0.18 (0.13) | −0.08 (0.48) | −0.05 (0.68) | −0.11 (0.36) |
| γ | −0.28 (0.02) | −0.19 (0.11) | −0.11 (0.34) | −0.19 (0.11) |

**Note.** Values represent Pearson's correlation coefficients (*r*) with corresponding uncorrected *p* values in parentheses. Correlations were calculated across participants (*n* = 74) between training-related changes (Training − Nontraining) in oscillatory power during the neurofeedback period and corresponding changes in pain intensity.

**Table S7.** Trial-level within-participant associations between target-S1 α power during neurofeedback and subsequent pain intensity.

| **Fixed effect** | **β** | **SE** | ***t*** | ***p*** |
|---|---|---|---|---|
| Feedback authenticity (Real vs. Sham) | −0.61 | 0.41 | −1.49 | 0.14 |
| Training (Training vs. Nontraining) | −0.90 | 0.08 | −11.94 | < 0.001 |
| Within-participant α | −0.15 | 0.05 | −2.75 | 0.006 |
| Training × Feedback authenticity | −0.72 | 0.10 | −6.90 | < 0.001 |
| α × Training | 0.15 | 0.07 | 1.97 | 0.049 |
| α × Feedback authenticity | −0.004 | 0.07 | −0.06 | 0.96 |
| α × Training × Feedback authenticity | −0.05 | 0.10 | −0.50 | 0.62 |

**Note.** Trial-level target-S1 α power, averaged across C4, CP4, and CP6, was quantified during the 16-s neurofeedback period and centered within each participant and experimental condition to capture within-participant fluctuations. Pain intensity was modeled using a linear mixed-effects model with centered α power, Training, Feedback authenticity, and their interactions as fixed effects and participant-specific random intercepts. Sham feedback and nontraining served as reference levels. The coefficient for α power represents its association with subsequent pain intensity in the sham-feedback/nontraining reference condition. β, regression coefficient; SE, standard error.

**Table S8.** Multiband trial-level mixed-effects model of target-S1 oscillatory power predicting subsequent pain intensity.

| **Frequency band** | **Power** | **Power × Training** | **Power × Feedback authenticity** | **Power × Training × Feedback authenticity** |
|---|---|---|---|---|
| δ | −0.10 (0.29) | −0.04 (0.74) | 0.23 (0.07) | −0.15 (0.40) |
| θ | 0.07 (0.48) | 0.04 (0.80) | 0.05 (0.69) | −0.25 (0.18) |
| α | −0.03 (0.69) | 0.02 (0.81) | −0.09 (0.34) | 0.04 (0.73) |
| β | −0.28 (0.02) | 0.25 (0.18) | 0.10 (0.59) | −0.01 (0.98) |
| γ | 0.19 (< 0.001) | −0.23 (0.002) | −0.03 (0.67) | 0.03 (0.75) |

**Note.** Trial-level oscillatory power across the five frequency bands (δ, θ, α, β, and γ), averaged across target-S1 electrodes (C4, CP4, and CP6), was centered within each participant and experimental condition and entered simultaneously into a linear mixed-effects model predicting subsequent pain intensity. The model included band-specific interactions with Training and Feedback authenticity, with participant-specific random intercepts. Sham feedback and nontraining served as reference levels. Values represent regression coefficients (β), with corresponding uncorrected *p* values in parentheses. The model also included the main effects of Training and Feedback authenticity and their interaction; the Training effect and Training × Feedback authenticity interaction were both significant (both *p* < 0.001).

## Notes

https://osf.io/x4gf2

### Competing Interest Statement

The authors have declared no competing interest.

### Summary of Updates

This version has been substantially revised through additional analyses, methodological clarifications, and refined interpretation of the findings. The major revisions include: 1. Expanded analyses of neural mechanisms and alternative explanations, including source-space analyses of alpha power and functional connectivity, examination of visual feedback characteristics, and sensitivity analyses of behavioral findings. 2. Improved characterization of EEG microstate dynamics, including a supplementary figure presenting the temporal characteristics of all five microstates and the complete set of directed transitions. The interpretation of the selective E-to-D transition effect has also been refined. 3. Additional analyses of the temporal dynamics of alpha modulation and its relationship with pain, examining alpha activity during neurofeedback and pre-pain periods, as well as trial-level and multiband associations between oscillatory activity and subsequent pain perception. 4. Improved methodological transparency and interpretation through clarification of methodological procedures, revision of figures and supplementary materials, and a more cautious discussion distinguishing feedback-dependent neural regulation from evidence linking preparatory neural changes to subsequent pain modulation. Exploratory structural equation modeling results are interpreted as indirect associations rather than established causal pathways. The accessibility of research materials has also been improved, and the completed CRED-nf checklist has been included to enhance transparency and reproducibility.

