## Supplementary material for "Closed-loop neurofeedback reshapes preparatory brain states to bias subsequent pain processing": CRED-nf Checklist

Checklist items are reproduced from Ros et al. (2020).

| **Domain** | **Item #** | **Checklist item** | **Reported on page #** |
| --- | --- | --- | --- |
| **Pre-experiment** | **1a** | Pre-register experimental protocol and planned analyses | Not preregistered. |
|  | **1b** | Justify sample size | p. 12 |
| **Control groups** | **2a** | Employ control group(s) or control condition(s) | p. 12 |
|  | **2b** | When leveraging experimental designs where a double-blind is possible, use a double-blind | p. 12 |
|  | **2c** | Blind those who rate the outcomes, and when possible, the statisticians involved | p. 12 |
|  | **2d** | Examine to what extent participants and experimenters remain blinded | pp. 4, 14 |
|  | **2e** | In clinical efficacy studies, employ a standard-of-care intervention group as a benchmark for improvement | N/A |
| **Control measures** | **3a** | Collect data on psychosocial factors | pp. 4, 12, 14 |
|  | **3b** | Report whether participants were provided with a strategy | p. 13 |
|  | **3c** | Report the strategies participants used | p. 4 |
|  | **3d** | Report methods used for online-data processing and artefact correction | pp. 13–14 |
|  | **3e** | Report condition and group effects for artefacts | Not reported. |
| **Feedback specifications** | **4a** | Report how the online-feature extraction was defined | pp. 13–14 |
|  | **4b** | Report and justify the reinforcement schedule | pp. 13–14 |
|  | **4c** | Report the feedback modality and content | pp. 13–14 |
|  | **4d** | Collect and report all brain activity variable(s) and/or contrasts used for feedback, as displayed to experimental participants | pp. 13–14 |
|  | **4e** | Report the hardware and software used | p. 13 |
| **Outcome measures — Brain** | **5a** | Report neurofeedback regulation success based on the feedback signal | p. 5 |
|  | **5b** | Plot within-session and between-session regulation blocks of feedback variable(s), as well as pre-to-post resting baselines or contrasts | pp. 5, 7 |
|  | **5c** | Statistically compare the experimental condition/group to the control condition(s)/group(s) (not only each group to baseline measures) | pp. 4–8 and 17 |
| **Outcome measures — Behaviour** | **6a** | Include measures of clinical or behavioural significance, defined a priori, and describe whether they were reached | pp. 4–8 and 17–18 |
|  | **6b** | Run correlational analyses between regulation success and behavioural outcomes | pp. 6–8 and 17 |
| **Data storage** | **7a** | Upload all materials, analysis scripts, code, and raw data used for analyses, as well as final values, to an open access data repository, when feasible | p. 18 |
